# MITF reprograms the extracellular matrix and focal adhesion in melanoma

**DOI:** 10.1101/2020.07.14.202291

**Authors:** Ramile Dilshat, Valerie Fock, Colin Kenny, Ilse Gerritsen, Romain Maurice Jacques Lasseur, Jana Travnickova, Ossia Eichhoff, Philipp Cerny, Katrin Möller, Sara Sigurbjörnsdóttir, Kritika Kirty, Berglind Ósk Einarsdottir, Phil F. Cheng, Mitchell Levesque, Robert Cornell, E. Elizabeth Patton, Lionel Larue, Marie de Tayrac, Erna Magnúsdóttir, Margrét H. Ogmundsdottir, Eiríkur Steingrímsson

**Author notes:** Institute of Molecular Life Sciences, University of Zurich, Y13-K-36, Winterthurerstrasse 190, CH8057 Zurich, Switzerland (Katrin Möller). Corresponding author: Eirikur Steingrimsson.

## Abstract

The microphthalmia associated transcription factor (MITF) is a critical regulator of melanocyte development and differentiation. It also plays an important role in melanoma where it has been described as a molecular rheostat that, depending on activity levels, allows reversible switching between different cellular states. Here we show that MITF directly represses the expression of genes associated with the extracellular matrix (ECM) and focal adhesion pathways in human melanoma cells as well as of regulators of epithelial to mesenchymal transition (EMT) such as CDH2, thus affecting cell morphology and cell-matrix interactions. Importantly, we show that these effects of MITF are reversible, as expected from the rheostat model. The number of focal adhesion points increased upon MITF knockdown, a feature observed in drug resistant melanomas. Cells lacking MITF are similar to the cells of minimal residual disease observed in both human and zebrafish melanomas. Our results suggest that MITF plays a critical role as a repressor of gene expression and is actively involved in shaping the microenvironment of melanoma cells in a cell-autonomous manner.

## Introduction

Melanoma is a highly aggressive form of skin cancer that originates from melanocytes. Approximately 60% of melanoma tumours harbour a BRAF mutation, most often BRAF_V600E_, which leads to hyperactivation of the mitogen-activated protein kinase (MAPK) pathway [1. Drugs targeting the BRAF and MAPK pathways are clinically important, but almost invariably, resistance arises within a short time period [2. Melanoma inherits its aggressive nature from its multipotent neural crest precursors that gives rise to various cells including melanocytes, glia and adrenal cells [3, 4]. The developmental programme of neural crest cells is believed to be reinitiated during melanoma progression and dysregulation of neural crest genes is predictive of metastatic potential and negative prognosis in melanoma [5–7]. Various different studies, including gene expression studies of tumours, immunohistochemical analysis of melanoma samples and single-cell sequencing studies of patient-derived xenografts suggest the existence of different cell types in melanoma tumours. This cellular heterogeneity is believed to reflect the associated ability of tumour cells to switch their phenotype from proliferative, non-invasive cells to quiescent, invasive cells and back, thus allowing metastasis and the escape from therapeutic intervention (reviewed in [8. This has been summarized in the phenotype switching model which suggests that melanoma cells can switch between invasive and proliferative states allowing them to either grow and form tumours or metastasize to a new site [8, 9]. Understanding the molecular mechanisms underlying the phenotypic plasticity of melanoma cells is key to addressing the metastatic potential of melanoma cells.

The Microphthalmia-associated transcription factor (MITF) is essential for melanocyte differentiation, proliferation and survival. MITF is also important during melanomagenesis (reviewed in [10). This is best evidenced by the observations that the rare germline mutation E318K of MITF increases the susceptibility to melanoma and MITF has been shown to be amplified in 15% of melanoma tumours [11–13]. Importantly, MITF activity has been used as a proxy for the phenotype switching model with MITFhigh cells characterized as proliferative whereas MITF_low_ cells have been assigned a quiescent invasive phenotype [14, 15]. In fact, MITF has been proposed to act as a rheostat where the levels of MITF activity determine the phenotypic state of melanoma cells (reviewed in [8). Since MITF expression and activity are regulated by the various signalling pathways, the tumour microenvironment has been proposed to instruct phenotypic changes in melanoma cells and thus foster disease progression [16–19]. However, antibody staining suggests that cells lacking MITF are abundant in melanomas [20 and single-cell sequencing of human xenotransplants and of zebrafish melanoma models suggest the existence of cells with very low MITF expression [21, 22]. These cells belong to a population of cells believed to represent minimal residual disease, cells that remain viable upon drug exposure.

The extracellular matrix (ECM) is an important component of the tumour microenvironment as it provides cells with biochemical and structural support. In melanoma, expression of ECM proteins such as tenascin and fibronectin increases during disease progression [23. Focal adhesions not only offer physical attachment of cells to the ECM through the integrin receptor, but also initiate signalling cascades that regulate cell proliferation, migration and survival [24–26]. A key focal adhesion signalling protein is Focal Adhesion Kinase (FAK), which activates the ERK pathway via Grb-FAK interactions [27. An important scaffolding protein at the focal adhesion complex is Paxillin (PXN) which recruits other proteins to the focal adhesion sites when phosphorylated by FAK and SRC [28. Importantly, phosphorylation of PXN is critical for activation of RAF, MEK, and ERK and has been shown to confer drug resistance by activating Bcl-2 through ERK signalling [29–33]. This highlights the importance of identifying a molecular mechanism that confers cells with the ability to circumvent drug inhibition through phenotypic changes.

In this study, we show that MITF represses the expression of focal adhesion and ECM genes in melanoma cells and tissues. Our findings reveal a new role for MITF in regulating the expression of genes that are essential for creating the melanoma microenvironment, establishing a link to melanoma progression and drug resistance.

## Results

### Melanoma cells devoid of MITF are enlarged and exhibit altered matrix interactions

To assess the effects of permanent loss of MITF in melanoma cells, we used the clustered regularly interspaced short palindromic repeats (CRISPR)-Cas9 technique to generate MITF knockout (KO) cell lines in the human hypo-tetraploid SkMel28 melanoma cell line (containing four copies of MITF). We targeted exons 2 (an early exon containing a transactivation domain) and 6 (containing the DNA binding domain) of MITF separately and the resulting isogenic cell lines are hereafter referred to as ΔMITF-X2 and ΔMITF-X6 (Fig. 1a). The control cell line EV-SkMel28 was generated by transfecting SkMel28 cells with Cas9 along with the empty gRNA plasmid. To identify mutations introduced in the cell lines, we performed whole genome sequence (WGS) analysis, which showed that mutations were introduced in MITF in both the ΔMITF-X2 and ΔMITF-X6 cells (Fig. 1b, c) but not in the EV-SkMel28 control. In addition, we confirmed the WGS analysis by amplifying the mutated genomic regions, cloning them into vectors and performing Sanger sequencing. The ΔMITF-X2 line had two different but independent insertion mutations in the same codon (insertion of A and T in the codon for Y22) and a 5-bp deletion (encoding Y22 and H23) that are present in 64%, 19% and 17% of sequenced DNA fragments in this region, respectively. All these mutations introduced frameshifts and premature stop codons in exon 2 of MITF (Fig. 1b). The mutations present in the ΔMITF-X6 line are the following: 52% of the sequenced fragments contained a deletion of 1-bp (encoding residue A198), 33% contained a 6 bp in-frame deletion in the basic domain of the protein (encoding residues R197-R198) and 15% of the sequenced fragments contained a 17-bp deletion (encoding residues 198-203). Both the 1- and 17-bp deletions introduced frameshifts and downstream stop codons (Fig. 1c) whereas the inframe 6-bp deletion removed two amino acids at the beginning of the alpha-helix encoding the basic domain and is therefore not expected to be able to bind DNA. No wild-type MITF gene was detected in either cell line. In both cell lines the ratio of mutants is consistent with two chromosomes carrying the same mutation and the remaining two chromosomes each carrying a different mutation. Western blotting revealed that the ΔMITF-X6 cells express very little, if any, MITF protein. Although the ΔMITF-X2 cells did not express the full-length ~55 kDa MITF protein, truncated forms of MITF were detected at ~40 and 47 kDa (Fig. 1d). These truncated forms were also present in wild type cells, albeit at lower levels, suggesting that these are alternative isoforms of the MITF protein (Fig. 1d). In order to determine if these shorter isoforms are due to alternative splicing, we performed RT-PCR across several exon-intron borders around exon 2 of the MITF transcript. Our results did not show any alternative splice forms of MITF (Supplementary Fig. 1a, b). The C5 MITF antibody used here recognizes an epitope located between residues 120 and 170 of MITF, which corresponds to exons 4 and 5 (Fig. 1a) [34. The truncated proteins observed in wild type and ΔMITF-X2 cells must still contain this region and are therefore likely to arise from alternative translation start sites. Immunostaining revealed a mostly nuclear staining of MITF in both the EV-SKmel28 and ΔMITF-X2 cells (Fig. 1e), indicating that the truncated MITF isoforms reside in the nucleus. However, in the ΔMITF-X6 cells, no signal for MITF was observed in the nucleus, whereas a very low background signal was observed in the cytoplasm (Fig. 1e). To summarize, we have generated two CRISPR MITF-KO cell lines from melanoma cells where ΔMITF-X6 is devoid of wild type MITF.

**Figure 1.**
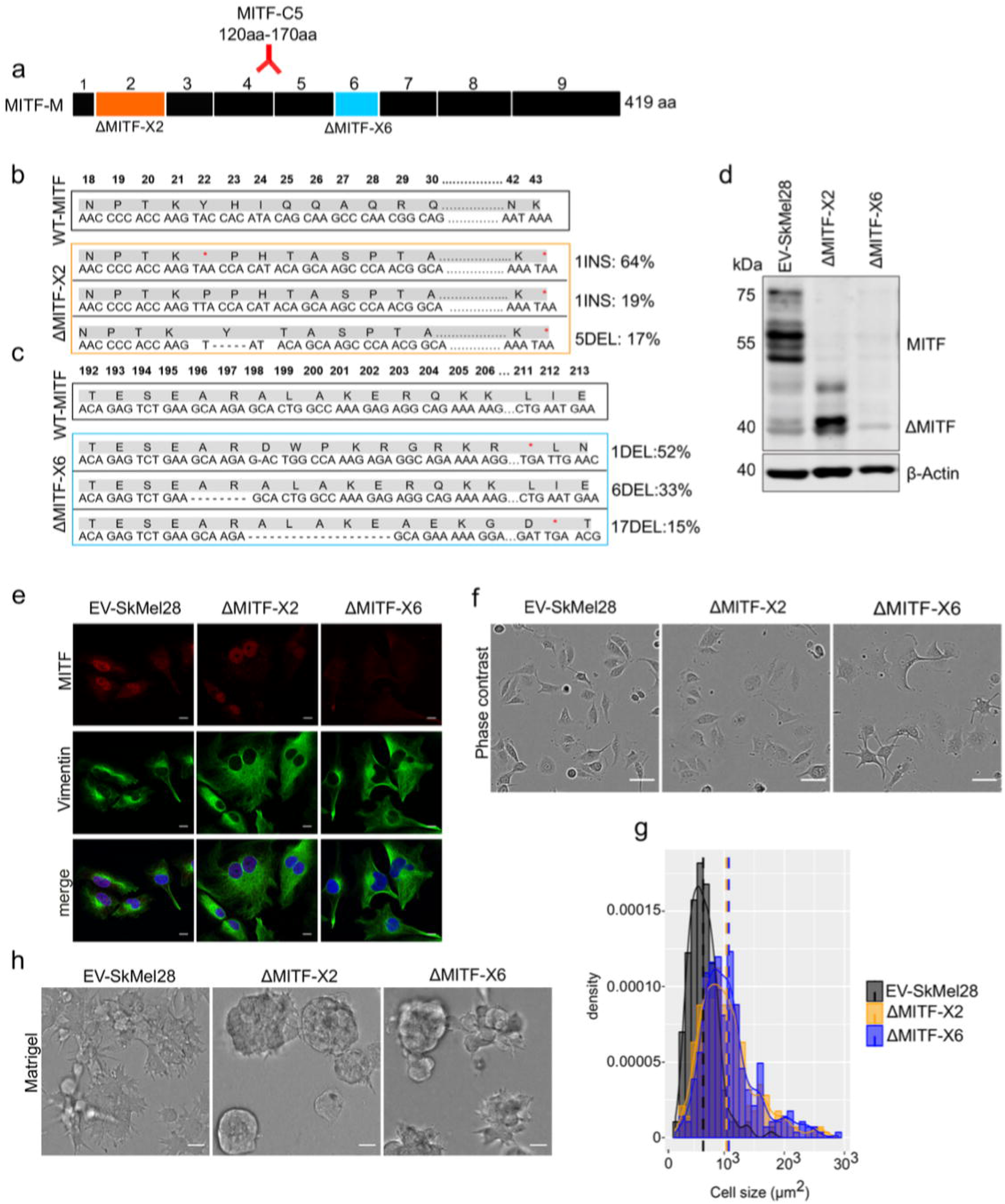
MITF depletion affects cell size and cell-matrix interaction. (a) Schematic illustration of MITF-M isoform and gRNA targeted location at exon 2 and exon 6. The epitope location for MITF C5 antibody spanning exon 4 and 5 is shown. (b, c) Mutations detected in ΔMITF-X2 and ΔMITF-X6 cell lines; amino acid sequence numbering was indexed relative to MITF-M. Percentage of mutations was derived from WGS analysis by counting sequenced fragments aligned to the mutated regions. (d) Western blot showing the MITF band in EV-SkMel28, ΔMITF-X2 and ΔMITF-X6 cell lines. (e) Immunostaining for MITF and Vimentin in EV-SkMel28, ΔMITF-X2 and ΔMITF-X6 cell lines, Scale bar (10μm). (f, g) Phase contrast microscopy and cell size quantification using Image J with at least 200 images taken for both MITF-KO and EV-SkMel28 cell lines, Scale bar (10μm). Average cell size for each cell line is indicated by dash line EV-SkMel28 cells (6,502 μm2, SEM: 460), ΔMITF-X2 (10,395 μm2, SEM: 270) and the ΔMITF-X6 (10,825 μm2, SEM: 330). (h) Bright field images of MITF-KO and EV-SkMel28 cells grown on top of matrigel, Scale bar (10μm).

Morphological analysis revealed that both MITF-KO cell lines exhibited enlarged cytoplasm as compared to controls (Fig. 1e-g). Vimentin staining revealed enlarged cells (Fig. 1e) which is consistent with a report showing that loss of MITF affects the cytoskeletal structure and shape of melanoma cells [14. Quantification of phase contrast microscopy images revealed that the average size of ΔMITF-X2 and the ΔMITF-X6 cells was 1.7-fold larger than the EV-SkMel28 cells (Fig. 1g). To characterize the behaviour of the cell lines when provided with ECM that mimics the basement membrane, we seeded the cells on top of matrigel-coated slides, supplemented with complete growth medium containing 2% matrigel. Both MITF-KO cell lines formed aggregates, whereas the control EV-SkMel28 cells displayed a flat sheet-like morphology (Fig. 1h). Taken together, our results show that loss of MITF lead to changes in cell morphology and cell-matrix interactions.

### Expression of ECM and focal adhesion genes is increased upon loss of MITF

Next we compared the transcriptomic profile of the ΔMITF-X6 cells (exhibiting complete loss of wild type MITF) to the EV-SkMel28 control cells. We identified 2,136 differentially expressed genes (DEGs) between ΔMITF-X6 and EV-SkMel28 cells with the cut off qval<0.05 (Supplementary Table 1). Of these, 1,516 genes showed 2-fold change in expression (Fig. 2a). Gene ontology and KEGG pathway enrichment analysis revealed that the genes reduced in expression upon MITF depletion were verified MITF-target genes involved in pigmentation and pigment cell differentiation such as *DCT, MLANA, OCA2* and *IRF4* in addition to *MITF* itself (Fig. 2a, b, Supplementary Table 1). Genes whose expression was increased upon loss of MITF were enriched in processes involved in glycosaminoglycan metabolism, extracellular matrix organization and extracellular structure organization, and included genes such as *SERPINA3, ITGA2, PXDN* and *TGFβ1* (Fig. 2a, b, Supplementary Table 1).

**Figure 2.**
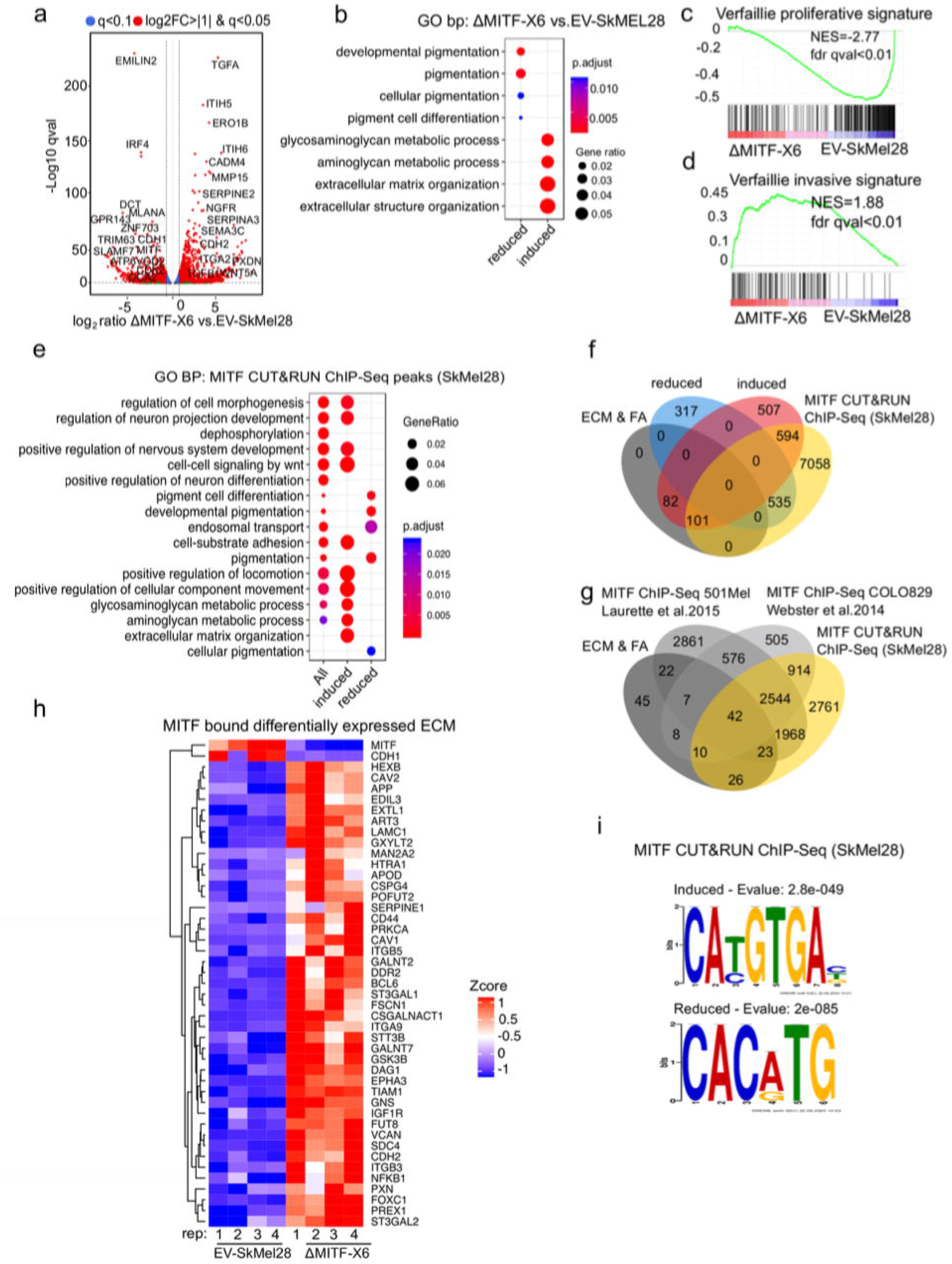
MITF binds and represses genes of ECM and focal adhesion genes. (a) Volcano plot showing 2,136 DEGs with qval<0.5 among which 1,516 genes with log2FC>|1| fold change in expression ΔMITF-X6 vs. EV-SkMel28 (b) GO BP analysis of DEGs 1,284 induced and 852 reduced between ΔMITF-X6 vs. EV-SkMel28 cells presented in dot plot; adjusted pvalue is red lowest to blue highest; gene ratio is the ratio between DEGs and all genes in the GO category. (c, d) Gene set enrichment analysis (GSEA) on pre-ranked DEGs of ΔMITF-X6 vs. EV-SkMel28 using Verfaillie [36 proliferative and invasive gene signature showing ΔMITF-X6 cells with positive enrichment for proliferative signature and negative enrichment for invasive signature. (e) Distribution of MITF CUT&RUN ChIP-Seq peaks in the genome. (f) GO BP analysis of MITF CUT&RUN ChIP-Seq peaks associated genes were plotted using Clusterprofiler [68 in R; Induced and reduced DEGs of ΔMITF-X6 vs. EV-SkMel28 cells based on MITF ChIP-seq peak presence on their gene promoter or distal region binding, and all MITF peak-associated genes regardless of DEGs. (g) Venn diagram showing the overlap between MITF targets identified from MITF CUT&RUN ChIP-Seq with induced, reduced, ECM and focal adhesion DEGs of ΔMITF-X6 vs. EV-SkMel28 cells (h) Venn diagram displaying the common overlap between MITF ChIP-Seq targets in different cell lines and differentially expressed ECM and focal adhesion genes in ΔMITF-X6 vs. EV-SkMel28 cells. (i) Heatmap showing the differentially expressed ECM genes in ΔMITF-X6 vs. EV-SkMel28 cells that are commonly bound by MITF across different MITF ChIP-Seq data sets. Zcore converted TPM value from RNA-Seq data was used for plotting. (j) Motif analysis of MITF CUT&RUN ChIP-Seq targets of induced and reduced genes in ΔMITF-X6 vs. EV-SkMel28 cells.

As MITF is central to the melanoma phenotype switching model [35, we were interested whether loss of MITF would be consistent with the published transcriptional signatures linked to phenotype switching in melanoma cells [36. Gene set enrichment analysis (GSEA) showed that the Verfaillie invasive signature displayed a positive enrichment with the ΔMITF-X6 cells, whereas the Verfaillie proliferative signature was negatively enriched (Fig. 2c, d). This further validated the MITF-KO cells as a representative model of long-term MITF loss.

In order to investigate if the genes affected by MITF loss are direct targets of MITF, we performed CUT&RUN MITF ChIP-Seq in the SkMel28 cells to map MITF genome wide binding sites. We identified 37,643 peaks located in the promoter, 3’UTR or intronic regions of 8,288 genes (Supplementary Fig. 2a) (Supplementary Table 2). Gene ontology analysis revealed that MITF-bound genes which showed increased expression upon MITF loss were enriched for aminoglycan, ECM and axogenesis pathways, whereas genes reduced in expression upon MITF loss were enriched for genes involved in pigmentation (Fig. 2e). We found that 695 of the 1,284 induced genes (P<7.3e-09 hypergeometric test) and 535 of the 852 repressed genes (P<6.6e-23, hypergeometric test) were directly bound by MITF (Fig. 2f) (Supplementary Table 2). Of the 183 ECM and focal adhesion genes whose expression was increased upon MITF knockout, 101 were bound by MITF and induced in expression upon loss of MITF (Supplementary Table 2). We compared our CUT&RUN MITF ChIP-seq peaks with the published MITF ChIP-Seq data from COLO829 (generated using the same antibody as used here) (Supplementary Table 3) and HA-MITF ChIP-Seq 501Mel cells (Supplementary Table 4) and found 42 ECM genes consistently bound by MITF in all three studies (Fig. 2g, h) (Supplementary Table 5).

To determine if the MITF peaks near induced and reduced genes contained the canonical MITF binding site, we performed *de novo* motif analysis of MITF-bound regions near DEGs using the MEMEChIP tool [37. We found that MITF-bound, induced genes (777) were primarily enriched for the 5’-CA[T/C]GTGAC-3’ motif, whereas MITF-bound reduced genes (535) were enriched for the 5’-CACATG-3’ motif (Fig. 2i). Thus, genes that are both induced and reduced in expression upon MITF loss contain MITF binding sites and are likely to be direct targets of MITF. Along with primary motifs, we also observed secondary motifs including a motifs for RUNX1and SOX10 in the induced, MITF-bound genes (Supplementary Fig. 2b). The secondary motifs observed in MITF-bound reduced genes were the FOXC1 like motifs (Supplementary Fig.2c). The differences observed in the secondary motifs may represent factors involved in repression versus activation functions of MITF. Taken together, we show that loss of MITF alters the expression of ECM genes and focal adhesions and large subset of them are directly bound by MITF.

### MITF depletion leads to increased expression of ECM genes

In order to verify that the link between MITF and the ECM and focal adhesion genes is not restricted to a particular cell line, we performed knockdown and overexpression studies in independent human melanoma cell lines and characterized gene expression data in the Cancer Genome Atlas. First, we performed mRNA sequencing after transient knockdown of MITF in SkMel28 and 501Mel cells, both of which express MITF endogenously at high levels. We identified 1,040 DEGs (qval<0.05, log2FC≥|1|, 567 induced, 473 reduced) upon siRNA-mediated MITF depletion in SkMel28 cells compared to siCTRL and 1,114 DEGs in 501Mel cells (qval<0.05, log2FC≥|1|, 624 induced, 490 reduced) (Supplementary Table 1). A significant correlation was observed between the DEGs of ΔMITF-X6 vs. EV-Skmel28 cells and DEGs of siMITF vs. siCTRL in SkMel28 (Pearson correlation=0.66, *P*<2.2e-16) and 501Mel cells (Pearson correlation=0.57, *P*<2.2e-16) (Fig. 3a, b). Second, we used the Cancer Genome Atlas dataset to characterize differential gene expression and split the tumours into two groups: the tumours with the 10% highest (MITFhigh) and 10% lowest (MITF_low_) expression of MITF. By performing differential gene expression analysis between the two groups, we identified 2,655 DEGs (FDR<0.01, log2FC>|1|, 1,835 induced and 820 reduced) between MITF_low_ and MITF_high_ tumours (Supplementary Table 1). Interestingly, the DEGs observed when comparing the ΔMITF-X6 cells to the EV-SkMel28 cells and the DEGs observed upon comparing the MITF_low_ and MITF_high_ tumours were significantly correlated (R=0.76, p<2.2e-1016) (Fig. 3c). Additionally, principal component analysis of the top 200 most statistically significant genes in each case revealed that MITF_low_ tumours cluster near the ΔMITF-X6 cells, whereas MITF_high_ tumours cluster with EV-SkMEL28 cells, indicating that ΔMITF-X6 cells portray the transcriptional state of MITF_low_ tumours (Fig. 3e). Third, we investigated whether overexpression of MITF would lead to repression of ECM genes. To do this, we performed mRNA-sequencing in A375P cells overexpressing a dox-inducible FLAG-tagged MITF construct (pB-MITF-FLAG). A control A375P cell line was generated using an empty vector only expressing FLAG (pB-FLAG). We identified 8,110 DEGs (qval<0.05, log2FC>| 11, 4,863 induced, 3,247 reduced) between pB-MITF-FLAG and pB-FLAG in A375P cells and among genes that are decreased in expression are ECM related genes (Supplementary Table 1). As expected, the DEGs observed upon MITF overexpression in A375P cells showed anti-correlation with the DEGs observed when comparing ΔMITF-X6 to EV-SkMel28 cells (Pearson correlation = −0.46, *P*<2.2e-16) (Fig. 3d).

**Figure 3.**
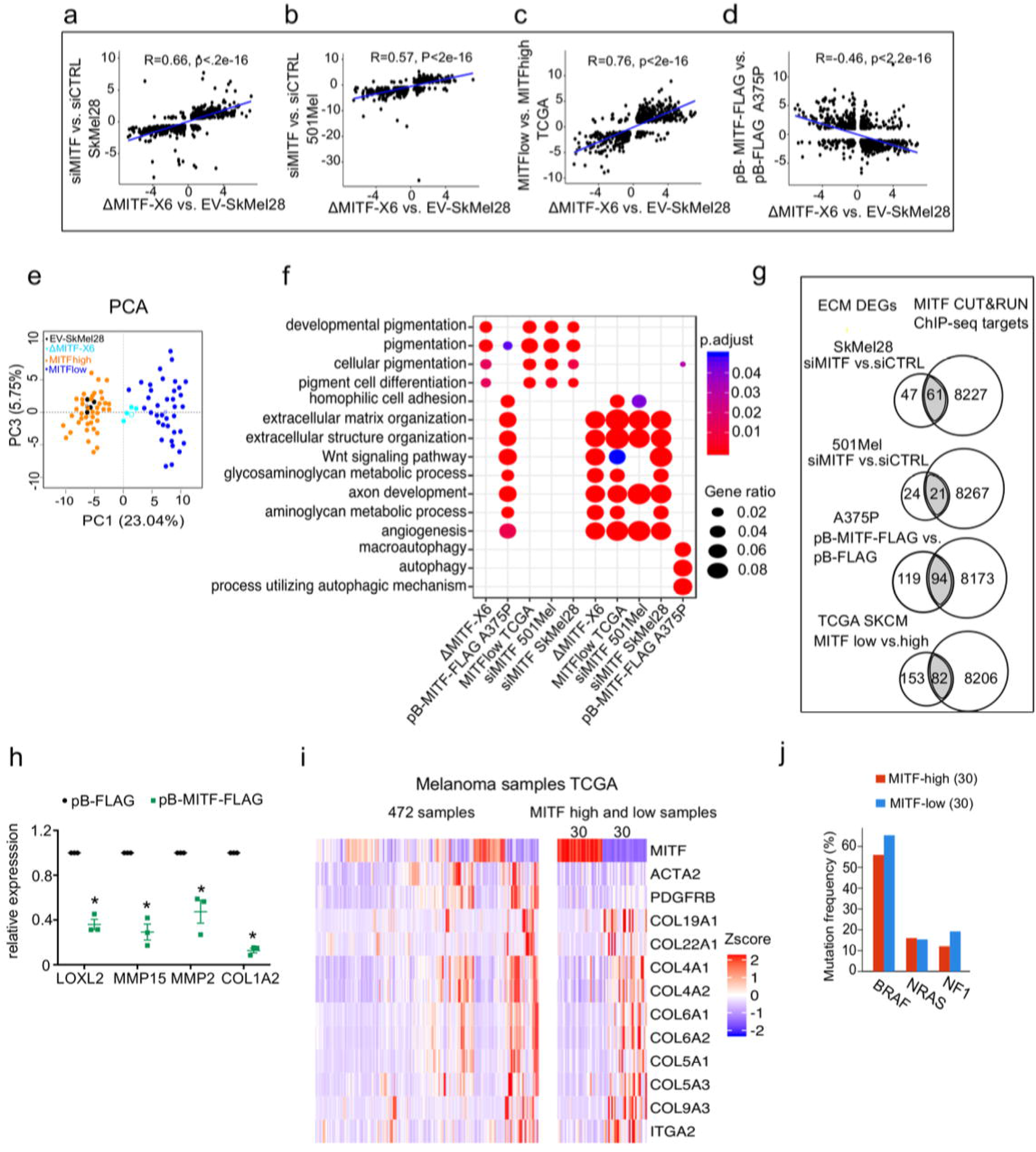
The ECM and focal adhesion gene signature is overrepresented upon MITF depletion and in MITF_low_ human melanoma tumours. (a-d) Positive correlation of DEGs in ΔMITF-X6 vs. EV-SkMel28 cells with DEGs of siMITF vs. siCTRL in 501Mel and SkMel28 and MITF_low_ vs. MITF_high_ melanoma tumours from TCGA, and negative correlation of DEGs in pB-FLAG vs. pB-MITF-FLAG A375P cells is shown. Values used in the X and Y axis are log2 fold change in the expression of DEGs. (e) Principal component analysis (PCA) of the most significant 200 DEGs in the MITF_low_ vs. MITF_high_ and ΔMITF-X6 vs. EV-SkMel28 display similar clustering of EV-SkMel28 samples with MITF_high_ tumours and ΔMITF-X6 cells with MITF_low_ tumours. (f) GO BP analysis of induced and reduced DEGs affected by MITF KO/KD in SkMel28 and 501Mel cells, and DEGs affected by MITF overexpression in A375P cells. (g) Venn diagram displaying the overlap in the number of differentially expressed ECM genes affected by MITF and MITF CUT&RUN ChIP-Seq targets. (h) RT-qPCR showing a reduced expression of ECM genes in the stable dox-inducible MITF overexpression A375P cell line (pB-MITF-FLAG). Relative expression was calculated by normalizing to control cells expressing empty vector (pB-FLAG). Error bars indicate standard error of the mean, (* pval < 0.05) was calculated using paired t-test. (i) Heatmap displaying the expression of ECM genes in the 472 melanoma samples from TCGA (left), from which the expression of ECM genes in the top 30 MITF_low_ and 30 MITF_high_ samples with high fibroblast marker removed (right) is shown. Hierarchical clustering was applied to cluster the samples Expression of genes was converted to Z-Score from red high to blue low. (j) Percentage of mutations in the MITF_low_ and MITF_high_ tumours from TCGA.

To classify genes that are overrepresented after the loss or gain of MITF we performed GO term enrichment analysis on the DEGs, which revealed an induction of ECM-related genes upon MITF depletion in 501Mel and SkMel28 cells as well as in MITF_low_ tumours, whereas genes involved in pigmentation were reduced in expression (Fig. 3f). Conversely, overexpressing MITF in the A375P cell line led to a reduction in expression of ECM genes and induction of pigmentation and autophagy genes, again showing that MITF negatively regulates the expression of ECM genes (Fig. 3f).

Analysis of the MITF ChIP-seq data [38 showed that a significant portion of the differentially expressed ECM genes upon MITF KD and in MITF_low_ tumours have MITF peaks in their regulatory domains (Supplementary Table 5) (Fig. 3g). In contrast, overexpression of MITF led to the repression of 213 ECM genes, 82 of which were MITF targets, indicating a major repressive influence of MITF on ECM gene expression (Fig. 3g) (Supplementary Table 5). We confirmed the repressive effects of MITF by RT-qPCR in dox-inducible A375P cells overexpressing pB-MITF-FLAG, which showed a significant reduction in the expression of *LOXL2, MMP15, MMP2,* and *COL1A2* when compared to control pB-FLAG cells (Fig. 3h). Together, our data support our conclusion that MITF is an important direct repressor of ECM gene expression in human melanoma cells and tissues.

Next, we analysed whether the collagens that were differentially expressed in the MITF-KD or KO melanoma cell lines were also affected by MITF in melanoma tumours in TCGA. Interestingly, we observed increased expression of collagen genes in the MITF_low_ tumours (Fig. 3i). However, to rule out the possibility that the increased expression of ECM genes in TCGA MITF_low_ melanoma tumours was derived from fibroblast cells, we removed the 130 melanoma TCGA samples that showed the highest expression of the fibroblast markers *PDGFRB* and *ACTA2* and then assessed the expression of collagens across the 30 MITF highest and lowest melanoma samples, which consistently showed that expression of genes encoding collagens are among the highest expressed genes in MITF_low_ tumours (Fig. 3i). We did not observe a correlation between MITF expression and the most common BRAF, NRAS and NF1 mutations found in melanoma (Fig. 3j), indicating that the gene expression changes observed are controlled via transcriptional regulation, directly or indirectly imposed by MITF. We conclude that reduced MITF expression leads to activation of expression of genes involved in ECM and focal adhesion in melanoma cells and tumours and that in many cases this is through direct binding of MITF to their regulatory regions.

### EMT genes are directly regulated by MITF

Genes involved in the EMT process have been shown to play a role in melanoma drug resistance and have been linked to low MITF expression [39, 40]. Consistent with this, analysis of the TCGA data showed that the expression of *CDH2 (N-cadherin), TGFB1* and *ZEB1* was anti-correlated with MITF in melanoma tumours whereas the expression of *CDH1 (E-cadherin)* and *SLUG (SNAI2)* was positively correlated (Fig. 4a). Consistent with this, the expression of the *CDH1* and SLUG genes was reduced in MITF_low_ tumours and ΔMITF-X6 cells whereas the expression of *CDH2*, *SOX2*, *TGFß1* and *ZEB1* was increased (Fig. 4b). We also observed increased expression of *CDH2* upon siRNA-mediated KD of MITF in SkMel28 and 501Mel cells, however the level of *CDH1* was decreased only in the siMITF SkMel28 cells (Fig. 4b). Interestingly, upon MITF overexpression in the pB-MITF-FLAG A375P cells, the expression of *CDH2, SNAI2, SOX2* and *TGFß1* was decreased whereas the expression of *CDH1* and *ZEB1* was increased (Fig. 4b). RT-qPCR analysis of EMT genes in the MITF-KO cells confirmed that *CDH1* expression was reduced 50- and 100-fold in the ΔMITF-X2 and ΔMITF-X6 cells, respectively, whereas *CDH2* and *TGFß1* were significantly increased when compared to EV-SkMel28 cells (Fig. 4c). Western blot analysis confirmed increased expression of the classical EMT marker protein CDH2 and decreased expression of CDH1 in both MITF-KO cell lines (Fig. 4d, e). Analysis of CUT&RUN ChIP-Seq and publicly available MITF ChIP-Seq data showed that *ZEB1, SOX2, CDH1* and *CDH2* genes contain MITF binding peaks in their intronic and promoter regions (Supplementary Fig. 2d) [38, whereas *TGFB1* does not. This suggests that MITF is not only involved in regulating the expression of ECM genes but may also be directly involved in regulating the expression of EMT genes, resulting in EMT-like changes in cell morphology and behaviour.

**Figure 4.**
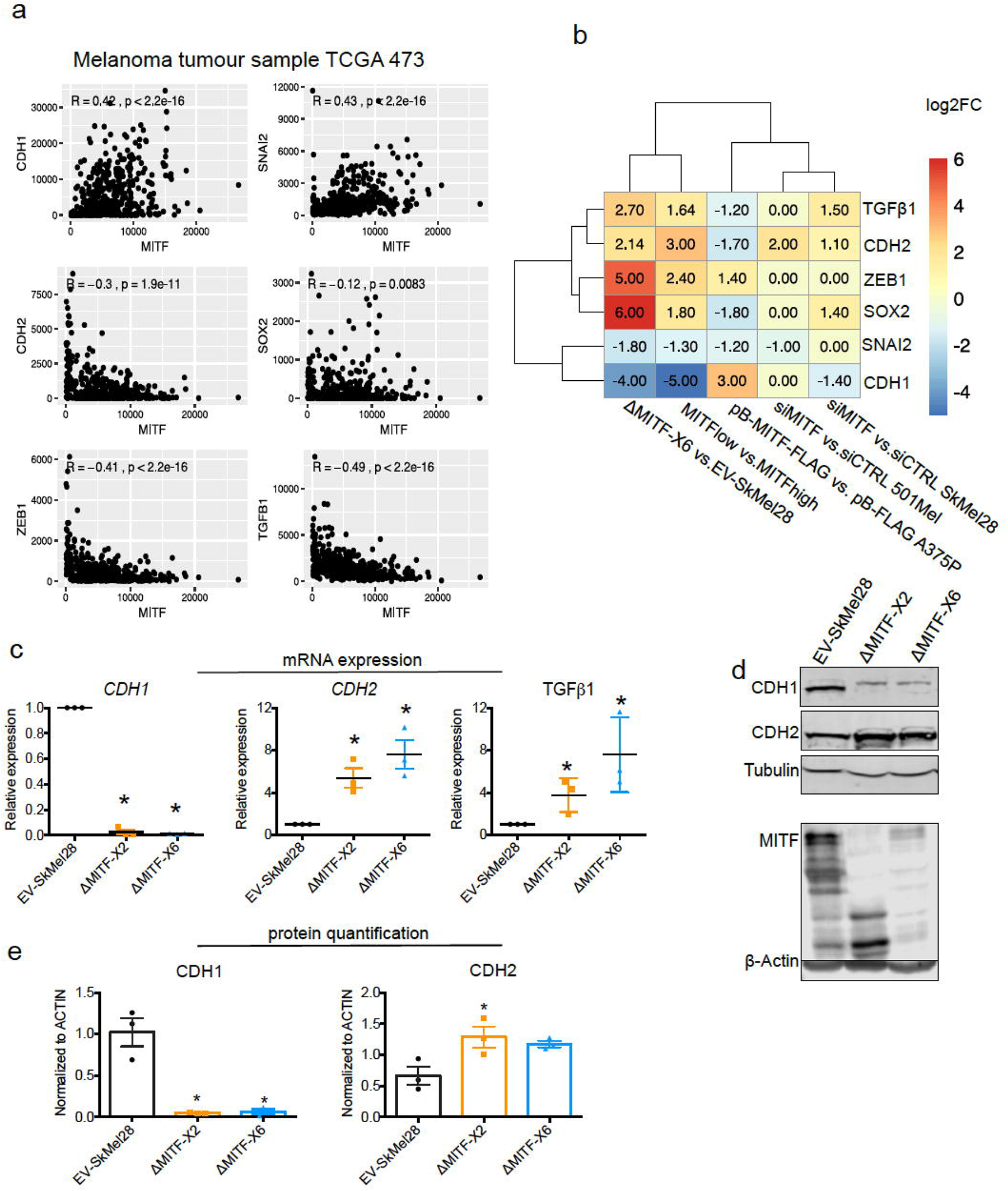
EMT genes are directly regulated by MITF. (a) Scatter plot displaying the spearman correlation between MITF mRNA expression with EMT genes in the 472 melanoma tumour samples from TCGA; MITF displayed positive correlation with *CDH1* and *SNAI2* and negative correlation with *ZEB1*, *TGFß1* and *CDH2*. (b) Differentially expressed EMT genes plotted as heatmap using the log2 fold change value of DEGs of MITF depletion in SkMel28 and 501Mel cells, MITF overexpression in A375P cells and MITF_low&high_ melanoma tumours. (c) Real-Time qPCR (RT-qPCR) evaluation of EMT genes in the EV-SkMel28, ΔMITF-X2 and ΔMITF-X6 cell lines. Fold change in the expression calculated over EV-SkMel28. Error bar represents standard error of the mean, (* pval < 0.05) was calculated using one-way ANOVA (multiple correction with Dunnett test). (d, e) Western blot analysis and quantification (Fiji Image J) of protein expression of CDH1, CDH2 and MITF in EV-SkMel28, ΔMITF-X2 and ΔMITF-X6 cell lines. ß-Actin was used as loading control. * pval < 0.05 was calculated by one-way ANOVA (multiple correction with Dunnett test).

### MITF-mediated effects on ECM genes are reversible

The MITF rheostat model predicts that different levels of MITF activity modulate distinct phenotypic states of melanoma cells and that these effects are reversible [41. To determine if the effects of long term MITF knockout could be reversed, we performed a rescue experiment by introducing an exogenous MITF-FLAG or EV-FLAG construct into the MITF-KO cells and then used RT-qPCR to characterize the expression pattern of ECM genes. As expected, the control EV-FLAG transfected MITF-KO cells exhibited increased expression of the ECM genes *CDH2, ID1* and *MMP15* as compared to the EV-SkMel28 control cells (Fig. 5a-d), whereas the expression of *CDH1* was reduced. Importantly, the expression of all four genes was partially rescued upon introducing the MITF-FLAG construct into ΔMITF-X6 cells; a smaller rescue effect was observed in ΔMITF-X2 cells transfected with MITF-FLAG (Fig. 5a-d).

**Figure 5.**
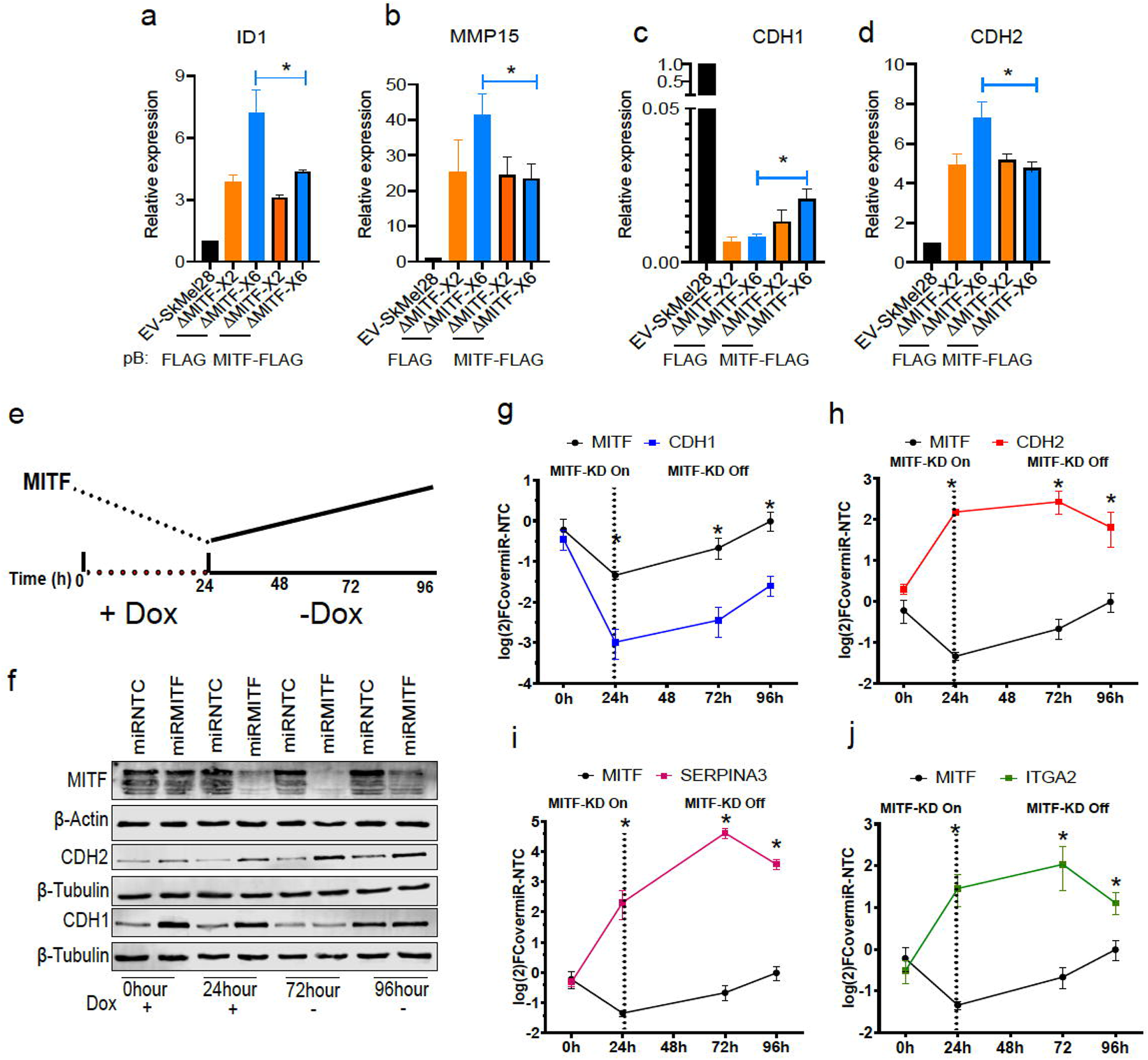
The effects of MITF on EMT and ECM gene expression are reversible. (a-d) Gene expression of ECM and EMT genes evaluated by RT-qPCR in EV-SkMel28 and MITF-KO cells with ectopic expression of EV-FLAG and MITF-FLAG constructs. Expression was normalised to EV-SkMel28 cells. Error bars represent standard error of the mean, * pval < 0.05 was calculated by two-way ANOVA (multiple correction with Sidak test). (e) Schematic showing the dox-inducible MITF KD system. MITF expression decreases in the presence of dox (first 24h) and reverts back to baseline levels upon dox wash-off (at 72h-96h). (f) Western blot analysis for the protein expression of MITF and CDH1, and CDH2 with the presence of dox treatment 0 and 24 hours or absence of dox 72 and 96 hours. (g-j) RT-qPCR analysis of MITF target ECM genes in miR-NTC and miR-MITF SkMel28 cells, treated with dox for 24h to induce MITF-KD and after dox wash-off at 72h and 96h. Expression was normalised to miR-NTC cell lines, error bars represent standard error of the mean, * pval < 0.05 was calculated by two-way ANOVA (multiple correction with Sidak test).

In order to overcome the partial rescue seen with the MITF-KO cells, we used the piggybac transposon system to integrate a dox-inducible synthetic micro-RNA construct (*miR-MITF*) into 501Mel and SkMel28 cells, thus allowing inducible knockdown (KD) of MITF by addition of doxycycline (Fig. 5e). At the same time, cells carrying a non-targeting control (*miR-NTC)* were generated. We induced MITF-KD in the miR-MITF SkMel28 cell line by adding dox and removed it again after 24 hours to assay for gene and protein expression at defined time points (Fig. 5e). We chose to focus on the ECM and EMT genes *CDH1, CDH2, ITGA2* and *SERPINA3,* all of which are direct targets of MITF (Supplementary Fig. 3) [38. Our results showed that MITF mRNA and protein expression was significantly decreased after 24 hours of dox treatment and reached basal levels again 96 hours after dox removal (Fig. 5f, g), showing that the dox-inducible system is suitable for reversibly modulating MITF levels. We observed a sharp decrease in *CDH1* mRNA expression after 24 hours of dox treatment. However, 72 and 96 hours after dox removal its expression had gradually increased, consistent with the restoration of *MITF* expression (Fig. 5g). Similarly, the expression of genes repressed by MITF such as *CDH2, SERPINA3,* and *ITGA2* was sharply increased after 24 hours of dox treatment and decreased again 96 hours after dox removal (Fig. 5h-j). Western blotting showed that the expression of the E-cadherin (CDH1) protein was reduced, whereas the expression of N-Cadherin (CDH2) was increased when compared to the *miR-NTC* control (Fig. 5f). After 72 and 96 hours of dox removal, MITF expression was restored and expression of the E-Cadherin protein was increased back to initial levels, whereas the expression of N-Cadherin was reduced compared to that observed at 24 hours of MITF-KD (Fig. 5f). These data show that, consistent with the rheostat model, the function of MITF as both a repressor and activator of gene expression has reversible effects on the expression of EMT and ECM genes.

### MITF affects the number of focal adhesions

Based on the observed increase in the expression of ECM and focal adhesion genes, we expected focal adhesion formation to be affected in the MITF-depleted cells. Indeed, immunostaining revealed an increased number of paxillin (PXN)-positive focal points (stained using PXN phospho-Tyr118 antibodies) around the cell periphery of MITF-KO cells as compared to EV-SkMel28 control cells (Supplementary Fig. 3a). Quantification of the focal points showed around 2-fold increase in their numbers in both MITF-KO cell lines (Supplementary Fig. 3b). Transcriptomic data of the 473 melanoma tumour samples from TCGA showed a significant negative correlation between the expression of MITF and PXN in these samples (Supplementary Fig. 3c). We also assessed the expression of *PXN* in a panel of 163 patient-derived melanoma cells exhibiting different levels of MITF. This showed that expression of *PXN* was specifically induced in MITF_low_ melanoma cell lines and displayed a negative correlation with *MITF* expression (Supplementary Fig. 3d, e). In order to evaluate whether the formation of focal adhesions would be induced upon short-term MITF loss, we integrated the dox-inducible *miR-MITF* transgene into 501Mel and SkMel28 cells and detected focal adhesions using the PXN antibody. After a 24-hour induction of MITF-KD, a 2-fold increase was observed in the number of PXN-positive focal points at the cell borders when compared to the *miR-NTC* control cell lines (Supplemental Fig. 3f-h). Analysis of ChIP-seq data showed an MITF peak in intron 6 of *PXN* containing the CACGTG motif (Supplementary Fig. 3i). This indicates that MITF affects the formation of focal adhesions by directly regulating the expression of PXN, a key player in focal adhesion.

Previous studies have shown that adaptive resistance to the BRAF_V600E_ inhibitor vemurafenib leads to activation of focal adhesion and ECM-related pathways [42. Indeed, treating the cells with vemurafenib led to a decrease in MITF protein expression in EV-SkMel28 cells which is consistent with the literature. However, the expression of MITF in the 501Mel cell upon vemurafenib treatment was increased compared to a DMSO control (Supplementary Fig.4a, b). This raises the question of whether the effects observed on ECM and focal adhesion genes upon BRAF inhibition are mediated through MITF. To evaluate the effects of BRAF inhibition on focal adhesions, we treated MITF-KO and EV-SkMel28 cells with vemurafenib and stained for phospho-paxillin (Tyr118). Consistent with the observation above, the MITF-KO cells showed a 4-fold increase in the number of focal adhesions as compared to EV-SkMel28 cells under the control DMSO-treated conditions (Fig. 6a (upper panel), b). Treatment with vemurafenib resulted in a significant increase in the number of focal adhesions in the EV-SkMel28 cells but a further increase was also observed in the MITF-KO cells (Fig. 6a (lower panel), b). Consistent with this, knockdown of MITF induced through the miR-MITF construct in both 501Mel and SkMel28 cells led to an increased number of focal adhesions when compared to *miR-NTC* cells (Fig. 6c, d (upper panels), e, f). Treatment with vemurafenib further increased the number of focal adhesions in SkMel28 cells expressing miR-NTC or miR-MITF, but again, more focal points were observed in miR-MITF cells under these conditions (Fig. 6c, d (lower panels)). Importantly, vemurafenib treatment alone did not lead to an increase in focal adhesion formation in 501Mel cells expressing *miR-NTC* which is consistent with increased MITF protein expression upon vemurafenib treatment whereas a major increase in focal adhesions was observed upon MITF depletion in the miR-MITF cells (Fig. 6c (lower panel), e). These results suggest that the formation of focal adhesions upon vemurafenib treatment is in part dependent on changes in MITF expression. However, since a further increase is observed in upon vemurafenib treatment of the knockout and knockdown lines, other factors must also be involved.

**Figure 6.**
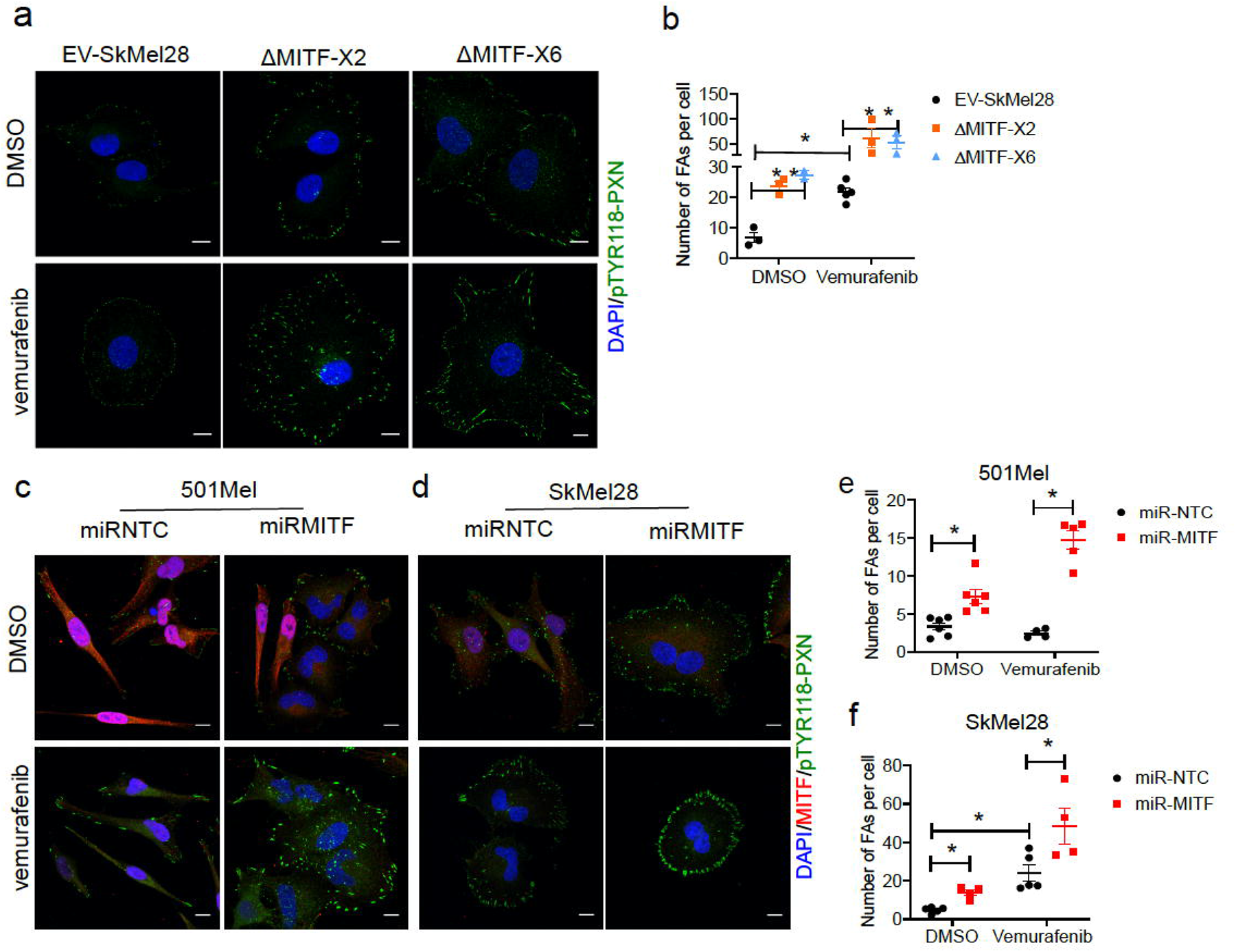
MITF mediates formation of focal adhesion. (a-c) Immunostaining for p-PXN_TYR118_ and quantification of p-PXN_TYR118_ positive focal points in EV-SkMel28, ΔMITF-X2 and ΔMITF-X6 cell lines treated with DMSO (a, upper panel) or vemurafenib (b, lower panel). (d-g) Immunostaining for p-PXN_TYR118_ and MITF and quantification of p-PXN_TYR118_ positive focal points in miR-NTC and miR-MITF 501Mel (d, f) and SkMel28 (e, g) cells. Error bars represent standard error of the mean, * pval < 0.05 was calculated by two-way ANOVA (multiple correction with Sidak test).

To understand whether the ECM and focal adhesion genes affected upon MITF loss overlap with the gene signature of melanoma cells that have been treated with BRAF inhibitors, we used single cell RNA-sequencing data of human melanoma xenografts [21. We focused on gene signatures specific for single cell populations with low MITF, (i) a subpopulation of cells which represent minimal residual disease (MRD) in melanoma, a small population of cells that remain upon drug treatment and (ii) an invasive gene signature [21. Our GSEA analysis showed that ΔMITF-X6 cells were significantly enriched in the MRD gene signature but not with the invasive signature found in another sub-population of MRD cells in the xenografts (Supplementary Fig. 4c). Among the genes that overlap between the MRD and ΔMITF-X6 cells are ECM genes such as *COL4A1, ITGA1, ITGA6, LAMC1* and *VCAN.* The same findings were obtained using single cell RNA-Seq data of MITF-depleted zebrafish melanomas as well as bulk-RNA-Seq data of MITF_low_ melanoma tumours [22. Both datasets showed positive enrichment with ΔMITF-X6 cells (Supplementary Fig. 4d). Importantly, we found that in the zebrafish data the ECM signature was specifically induced in the single cell cluster from MITF-low superficial tumours (representing minimal residual disease) compared to other single cell clusters from MITF-high melanomas (Supplementary Fig. 4e). These results suggest that the loss of MITF is an important mediator of MRD in melanoma and that MRD cells alter their extracellular environment.

### MITF KO affects proliferation and migration

The rheostat model predicts that MITF loss should reduce cell proliferation but increase migration potential of melanoma cells. We therefore measured proliferative ability of the MITF-KO cells using different methods. First, we characterized cell confluency over time using IncuCyte live cell imaging. This showed that both of the MITF-KO cells had a two-fold reduction in proliferation rate as compared to the EV-SkMel28 cells (Fig. 7a). Second, a BrdU incorporation assay showed that ΔMITF-X6 and ΔMITF-X2 cells had fewer (20%-25%) BrdU positive cells than the EV-SkMel28 (45%), suggesting that there are fewer actively proliferating cells in the MITF-KO cells compared to the control cells (Fig. 7b).

**Figure 7.**
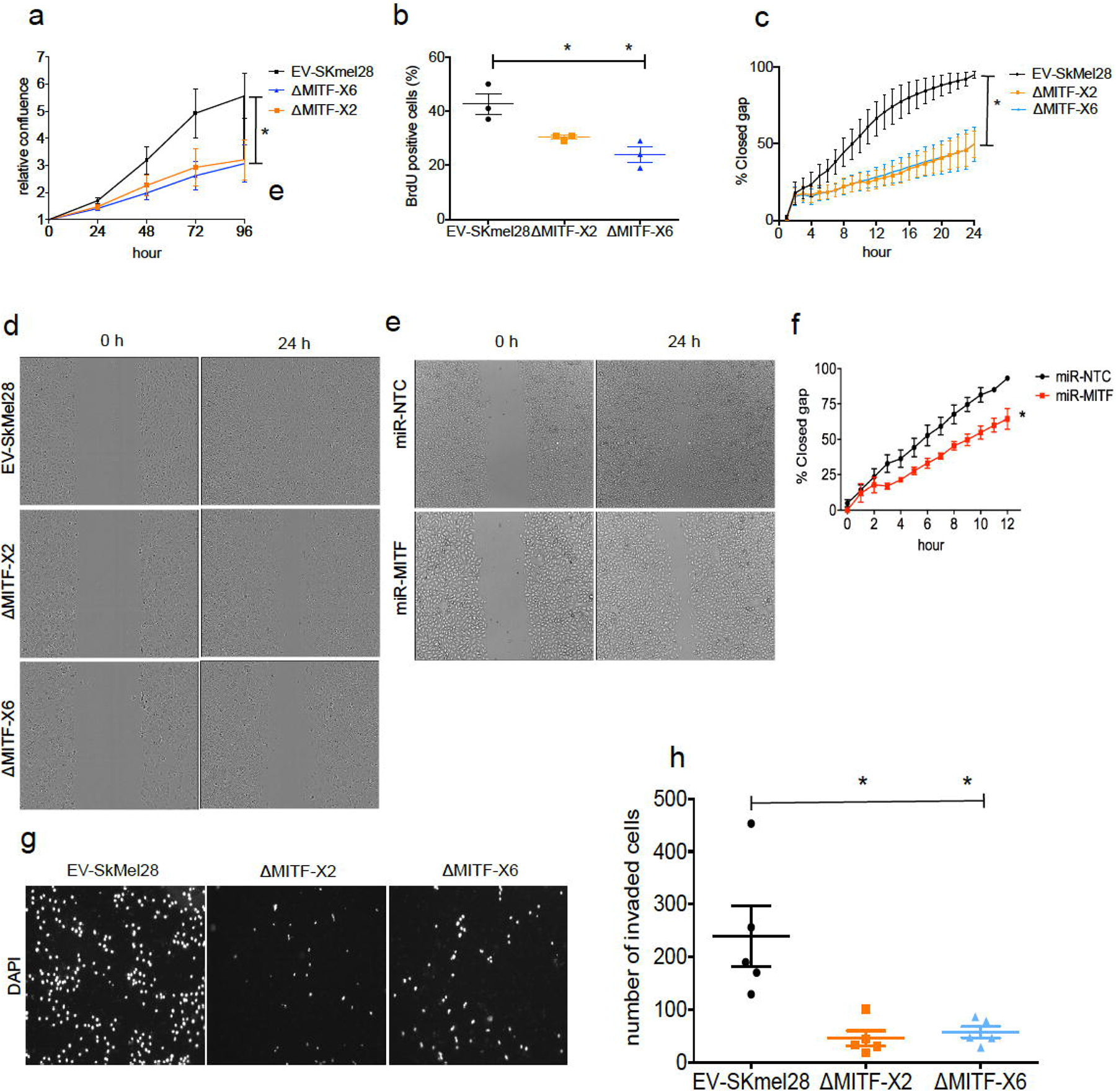
MITF knockout affects proliferation, migration and invasion ability of melanoma cells. (a) Relative cell confluency obtained from IncuCyte live cell imaging compared to day 0 was plotted for EV-SkMel28, ΔMITF-X2 and ΔMITF-X6 cell lines; Error bars represent standard error of the mean, * p-val < 0.05 was calculated by one-way ANOVA. (b) Percentage of BrdU positive cells was assessed by flow cytometry in EV-SkMel28, ΔMITF-X2 and ΔMITF-X6 cell lines. Error bars represent standard error of the mean, * pval < 0.05 was calculated by one-way ANOVA (multiple correction with Dunnett test). (c, d) Quantification and images of wound scratch assay in EV-SkMel28, ΔMITF-X2 and ΔMITF-X6 cells over 24 hour time period Error bars represent standard error of the mean, * p-val < 0.05 was calculated by two-way ANOVA (multiple correction with Sidak test). (e, f) Quantification and images of wound scratch assay in miR-NTC and miR-MITF in SkMel28 cells over 12 hour time period. Error bars represent standard error of the mean, * p-val < 0.05 was calculated by one-way ANOVA (multiple correction with Sidak test). (g, h) Matrigel invasion assay of EV-SkMel28, ΔMITF-X2 and ΔMITF-X6 cells using transwell; Quantification of invaded cells per transwell filter. Error bars represent standard error of the mean, * pval < 0.05 was calculated by one-way ANOVA (multiple correction with Dunnett test).

Previous analysis has shown that knocking down MITF leads to increased migration ability of melanoma cells [14, 43–47]. We therefore characterized the migration ability of our knockout cells. Strikingly, the wound scratch assay showed that the MITF-KO cells failed to close the wound in 24 hours whereas the EV-SkMel28 cells were able to close the wound within that time (Fig. 7c, d). To test whether the effects on migration were due to the long-term depletion of MITF in the MITF-KO cells, we performed the wound scratch assay upon MITF KD in the miR-MITF cells. Upon MITF-KD, we observed a minor decrease in the ability of the cells to close the wound when compared to the control miR-NTC cells (Fig. 7e, f). Next, we assessed the invasion ability of the MITF-KO cells using transwell chambers coated with matrigel. Interestingly, we found that MITF-KO cells displayed a severe reduction in invasion ability compared to the EV-SkMel28 cells (Fig. 7g,h). Taken together our data suggests that knocking down MITF negatively influences both cell proliferation and migration ability of the cells.

## Discussion

In this study, we have shown that MITF directly binds to and represses the expression of ECM, EMT and focal adhesion genes in human melanoma cells. We first observed this using our MITF-KO cells but verified our observations in other cell models by overexpression and knockdown of MITF using siRNA and inducible microRNA against MITF (*miR-MITF*) in melanoma cells. Importantly, we showed that MITF_low_ tumours in humans as well as in zebrafish have increased expression of ECM and focal adhesion genes. Together, our findings indicate that MITF acts as a transcriptional repressor of genes involved in ECM and focal adhesion.

A role for MITF as a repressor has been described in both melanoma cells and immune cells [47, 48]. In myeloid precursor cells, MITF was shown to interact with EOS to recruit co-repressors to target genes [48 whereas in melanoma cells MITF bound directly to an E-box located in an enhancer of the *c-JUN* gene, leading to reduced expression of the gene [19. Our results show that many of the genes whose expression is repressed by MITF are bound by MITF and contain E-boxes in their regulatory regions (Fig. 2i). This suggests that direct binding of MITF is involved in their repression. Since we observed differences in secondary motifs between the repressed and activated genes, different co-factors may be involved in mediating the repression in each case.

The MITF-dependent rheostat model which explains the phenotype switching of melanoma cells proposes that high MITF activity dictates proliferation whereas low MITF activity results in an invasive phenotype [14. Consistent with the rheostat model, proliferation was severely reduced upon MITF knockout (Fig. 7a, b). Unexpectedly, however, the migrative and invasive properties were reduced in both MITF-KO and MITF KD (*miR-MITF*) cells (Fig.7c-h). Immunohistochemistry and single-cell sequencing studies of melanoma tumours have shown the existence of cells with low or no MITF expression [20–22]. The involvement of MITF in migration has mostly been characterized using knockdown studies in melanoma cell lines using either siRNA or shRNA and by using Matrigel-coated Boyden chambers [14, 44–46]; in these studies knocking down MITF resulted in increased migration properties. Cheli et al. (2012) [44 also injected melanoma cells into the tail vein of mice and showed increased formation of metastasis when MITF was knocked down. Two different pathways involved in migration were shown to be regulated by MITF; DIAPH1, a gene implicated in actin polymerization [14, and the guanosine monophosphate reductase (GMPR) gene encoding an enzyme involved in regulating intracellular GTP levels [46. Surprisingly, however, more recent studies by Falletta et al. [47 and Vlckova et al. [49 failed to observe any effects on migratory/invasive properties upon MITF knockdown using the same cell lines as were used in the previous studies. Interestingly, knocking down SMAD7 in melanoma cells resulted in a dual invasive-proliferative phenotype without affecting MITF expression [50 Thus, the idea has been proposed that two different modes of invasion operate in melanoma; one with low MITF levels and no proliferation and another with high MITF levels where proliferation and invasion can take place simultaneously [10. Transcriptomic analysis displayed loss of expression of genes involved in proliferation, whereas invasive genes linked to AP1 and TEAD transcription factors was increased in the MITF-KO cells (Fig. 2c, d); the expression of TEAD and AP1 was not changed in our models. Thus, losing MITF alone likely does not lead to cells with invasive ability and something else is needed for gaining this property.

We identified MITF as an important transcriptional regulator of ECM and focal adhesion genes. Interestingly, we observed increased expression of TGFß1, an important regulator of ECM-related genes in the MITF-KO cells and MITF_low_ melanoma tumours (Fig. 4b, c). It has been shown that TGFß1 supresses the expression of MITF in melanoblasts, thereby inhibiting differentiation into melanocytes [51. This autocrine signalling of TGFβ is retained in melanoma cells [52. According to Hoek et al. (2006) [15, the MITF_low_ transcriptional state is dictated by TGFß1 signalling, which can suppress MITF expression resulting in an invasive and drug resistant phenotype [15, 17]. This suggests that the genes induced upon MITF loss are partly due to induction of TGFß signalling. However, our results suggest that MITF is directly involved in mediating the observed effects on the expression of ECM and focal adhesion genes. In addition, the relationship between MITF and the expression of TGFß1 is not clear. Our observations suggest that knocking down MITF leads to a major increase in TGFß1 mRNA expression in the melanoma cells, suggesting that the effects are cell-autonomous and driven by MITF. However, there are no MITF-peaks in or near the TGFß1 gene in melanoma cells, leading us to hypothesise that the effects must be mediated through a hitherto unknown intermediary.

Enhanced expression and phosphorylation of paxillin has been linked to therapy resistance in other cancer cell types, such as lung cancer [29. In melanoma, an inverse relation between BRAF inhibition and the expression of ECM genes has been described as a marker of de-differentiated drug resistant cells [42. Our data showed that the number of paxillin-positive dots was induced in both MITF-KO and miR-MITF cells as compared to controls (Supplementary Fig. 3a, b, f, g h) and paxillin expression was inversely correlated with MITF expression in melanoma tissues and cell lines (Supplementary Fig. 3c-e). Interestingly, we found that treating cells devoid of MITF with a BRAF inhibitor resulted in an increase in formation of focal adhesions (Fig. 6a-f). It is worth mentioning that an increase in the number of focal adhesion was restricted to SkMel28 melanoma cells in which MITF protein level was reduced upon vemurafenib treatment (Fig. 6d, f, Supplementary Fig.4a, b). However, we did not observe a significant increase in the number of focal adhesion in the 501Mel cells that gained MITF upon vemurafenib treatment (Fig.6 c, e, Supplementary Fig.4a, b). This highlights the role of MITF as a mediator of focal adhesion formation. However how the synergistic effects of MITF and vemurafenib on focal adhesion formation are mediated is unclear. One way to explain an increase in the formation of focal adhesions is that it is due to integrin clustering that is essential for the activation of focal adhesion pathways [53, 54]. We observed an increase in the expression of several integrins including *ITGA1, ITGA2, ITGA6, ITGA10* and *ITGB3* in the MITF-KO cells, as well as in the siMITF 501Mel and SkMel28 cell lines (Supplementary Table 5). In addition to this, the FLT1 receptor tyrosine kinase (VEGFR1) and its ligand VEGFA, which activate a pathway that phosphorylates FAK, a key mediator of focal adhesions, were increased in expression. Interestingly, both *FLT1* and *VEGFA* have MITF binding sites in their promoters and MITF has previously been shown to regulate *VEGFA* expression [55. Exposure of melanoma cells to BRAF and MEK inhibitors has been shown to slow growth and result in increased expression of *NGFR* and ECM and focal adhesion genes [42. Consistent with these findings, we observed an up to 200-fold induction of the *NGFR* transcript in the MITF-KO cells compared to EV-SkMel28 cells, and we identified an MITF peak in the 3’UTR of *NGFR* in both the MITF CUT&RUN ChIP-Seq data from SkMel28 cells and in the COLO829 cells [56; expression of the melanocyte differentiation marker and MITF target *MLANA* was 50-80 fold reduced in the MITF-KO cells (Supplementary Fig. 5a-d) Thus, it is possible that MITF affects focal adhesions by both directly regulating expression of genes involved in the process and indirectly by activating the expression of signalling processes involved.

Upon MITF loss, an EMT-like process has been described to be involved in driving drug resistance in melanoma [39, 40]. In addition, the degree of plasticity between EMT and mesenchymal to epithelial transition (MET) has been suggested to lead to high metastatic potential as well as therapeutic resistance [57–59]. Indeed, we observed changes in important EMT markers and regulators such as *ZEB1*, E-Cadherin, N-Cadherin, *Slug* and *TGFß1* in the MITF-KO cells (Fig. 4b-e) as well as in TCGA melanoma samples. Also, the MITF-KO cells showed increased expression of *SOX2,* which is important for neuronal stem cell maintenance and has been suggested to be important for self-renewal of melanoma tumour cells [60, 61] (Fig. 4b). Importantly, the effects of MITF on the expression of E-Cadherin, N-Cadherin and ECM genes (ITGA2 and SERPINA3) is reversible (Fig. 5e-j). This suggests that MITF enables epithelial to mesenchymal plasticity (EMP) that allows the formation of a hybrid state between EMT and MET to enforce the aggressiveness of melanoma. The binary effects of MITF on the expression of EMT genes may be the molecular mechanism that explains its rheostat activity.

The minimal residual disease (MRD) is a major reason for relapse in cancer. We found that ΔMITF-X6 cells are positively correlated with the gene signature of a population of MRD cells in melanoma tumours as determined by single-cell RNA-Seq of human PDX samples and zebrafish melanoma models (Supplementary Fig. 4c-e) [21. Interestingly, the MRD melanoma cells in zebrafish express little to no MITF protein and have increased expression of ECM genes (Supplementary Fig. 4e). This suggests that the induced expression of ECM genes and low expression of MITF is one of the markers of MRD in melanoma. In the absence of MITF, melanoma cells may become MRD cells by reshaping their extracellular matrix, enhancing their attachment to the surface, thus forming quiescent cells which wait for an opportunity to change their phenotype and re-emerge as proliferative melanoma cells. Since melanoma cells can mediate these effects on their own, in the absence of the tumour microenvironment, this suggests that this process is cell-autonomous and under the direction of MITF which instructs the cells to create their own microenvironment.

## Material and methods

### Cell culture, reagents and antibodies

SkMel28 cells were purchased from ATCC (HTB-72) and 501Mel melanoma cells were obtained from the lab of Ruth Halaban. The cells were grown in RPMI 1640 medium (#5240025, Gibco) supplemented with 10% FBS (#10270-106, Gibco) at 5% CO2 and 37°C. We made stocks of 5mM FAK inhibitor (Selleckchem, PF562271) and 5mM vemurafenib (Selleckchem, S1267) in DMSO and used a dilution of 1μM final concentration in cell culture media in all drug treatment experiments. The following primary antibodies and their respective dilutions were used in immunofluorescence (IF) and Western blot (WB) experiments: MITF (C5) mouse monoclonal (Abcam, #ab12039), 1:2000 (WB), 1:200 (IF); Phospho-Paxillin (Tyr118) rabbit monoclonal (Cell signalling, #2541), 1:000 (WB), 1:100 (IF); Vimentin rabbit monoclonal (Cell signalling, #3932), 1:100 (IF); ERK (p44/42 MAPK (Erk1/2), CST #9102) 1:1000 (WB); p-ERK (Phospho-p44/42 MAPK (Erk1/2) (Thr202/Tyr204) CST #9101) 1:1000 (WB); E-Cadherin (#610182, BD) 1:5000 (WB), N-Cadherin (#610921, BD) 1:5000 (WB); β-Actin rabbit monoclonal (Cell signalling, #4970), 1:2000 (WB), 1:200 (IF); β-Actin rabbit mouse monoclonal (Millipore, #MAB1501), 1:20000 (WB).

### Generation of MITF-KO cells and validation of mutations using Sanger sequencing

The CRISPR/Cas9 technology was used to generate knock out mutations in the MITF gene in SkMel28 cells. These cells carry the BRAFV600E and p53L145R mutations [62. Guide RNAs (gRNAs) were designed targeting exons 2 and 6 of MITF, both of which are common to all isoforms of MITF; exon 2 encodes a conserved domain of unknown function as well as a phosphorylation site, whereas exon 6 encodes the DNA binding domain of MITF (Fig. 1a). The gRNAs used were:AGTACCACATACAGCAAGCC (Exon2-gRNA); AGAGTCTGAAGCAAGAGCAC (Exon6-gRNA). The gRNAs were cloned into a gRNA expression vector (Addgene plasmid #43860) using BsmBI restriction digestion. The gRNA vectors were transfected into SkMel28 melanoma cells together with a Cas9 vector (a gift from Keith Joung) using the Fugene® HD transfection reagent (#E2312 from Promega) at a 1:2.8 ratio of DNA:Fugene. After transfection, the cells were cultured for 3 days in the presence of 3μg/ml Blasticidin S (Sigma, stock 2.5mg/ml) for selection and then serially diluted to generate single cell clones. As a result, we obtained the ΔMITF-X2 cell line from targeting exon 2 of MITF and the ΔMITF-X6 cell line from targeting exon 6. The respective control cell line, termed EV-SKmel28, was generated by transfecting the cells with empty vector Cas9 plasmid.

Genomic DNA was isolated from the MITF knock out cell lines using the following procedures: Cells (~ 2 x 105) were trypsinized and spun down and the supernatant was removed. The cell pellet was resuspended in 25 μL of PBS. Then 250 μL Tail buffer (50mM Tris pH8, 100 mM NaCI, 100 mM EDTA, 1% SDS) containing 2.5 μL of Proteinase K (stock 20 mg/mL) were added to the cell suspension in PBS and incubated at 56°C overnight. Then 50 μL of 5M NaCl were added and mixed on a shaker for 5 minutes and spun at full speed for 5 minutes at room temperature. The supernatant was then transferred into a new tube containing 300 μL isopropanol, mixed by inversion and spun in a microfuge for 5 minutes at full speed. The resulting pellet was washed with 70% ethanol and the pellets air-dried at room temperature. Finally, the dried pellets were dissolved in nuclease free water for at least 2 hours at 37 °C. The appropriate regions (exons 2 or 6) of MITF were amplified using region-specific primers (MITF-2-Fw: CGTTAGCACAGTGCCTGGTA, MITF-2-Rev: GGGACAAAGGCTGGTAAATG; MITFexon6-fw: GCTTTTGAAAACATGCAAGC, MITFexon6-rev: GGGGATCAATTCTCCCTCTT). The amplified DNA was run on a 1,5% agarose gel, at 70V for 60 minutes. The bands were cut out of the gel and extracted using Nucleospin Gel and PCR Cleanup Kit (#740609.50 from Macherey Nagel). The purified DNA fragments were cloned into the puc19 plasmid and 10 colonies were picked for each cell line, DNA isolated and sequenced using Sanger sequencing. Whole genome sequencing was performed using total genomic DNA isolated from the EV-SkMel28 as MITF-KO cells using the genomic isolation procedure above. Sequencing results were analysed using R package CrispRVariant [63 in Bioconductor to quantify mutations introduced in the MITF-KO cell lines.

### Generation of plasmids for stable doxycycline-inducible MITF knock down and overexpression cell lines

The piggy-bac transposon system was used to generate stable inducible MITF knockdown cell lines. The inducible promoter is a Tetracyclin-On system, which is called reverse tetracyline-transactivator (rtTA). This system allows the regulation of expression by adding tetracycline or doxycycline to the media. We used a piggy-bac transposase vector from Dr. Kazuhiro Murakami (Hokkaido University) [64. The microRNAs targeting MITF (Table 1) were cloned into the piggy-bac vector downstream of a tetracycline response element (TRE). First, we used BLOCK-iT RNAi designer to design microRNAs targeting MITF (exons 2 and 8 of MITF), including a terminal loop and incomplete sense targeting sequences that are required for the formation of stem loop structures (Table 1). To obtain short double-stranded DNAs with matching BsgI overhangs, the mature miRNAs were denatured at 95°C, then allowed to cool slowly in a water bath for annealing. Then the piggy-bac vector pPBhCMV1-miR(BsgI)-pA-3 was digested with BsgI (#R05559S, NEB) and the digested vector excised from a DNA agarose gel and the DNA purified. Following this, the annealed primers and purified digested vector were ligated at a 15:1 insert to backbone molar ratio using Instant Sticky-end Ligase Master Mix (M0370S, NEB). A non-targeting control (miR-NTC) was used as a negative control. The ligation products were then transformed to high-competent cells, clones isolated and plasmid DNA sequenced to verify the successful ligation.

**Table 1.**
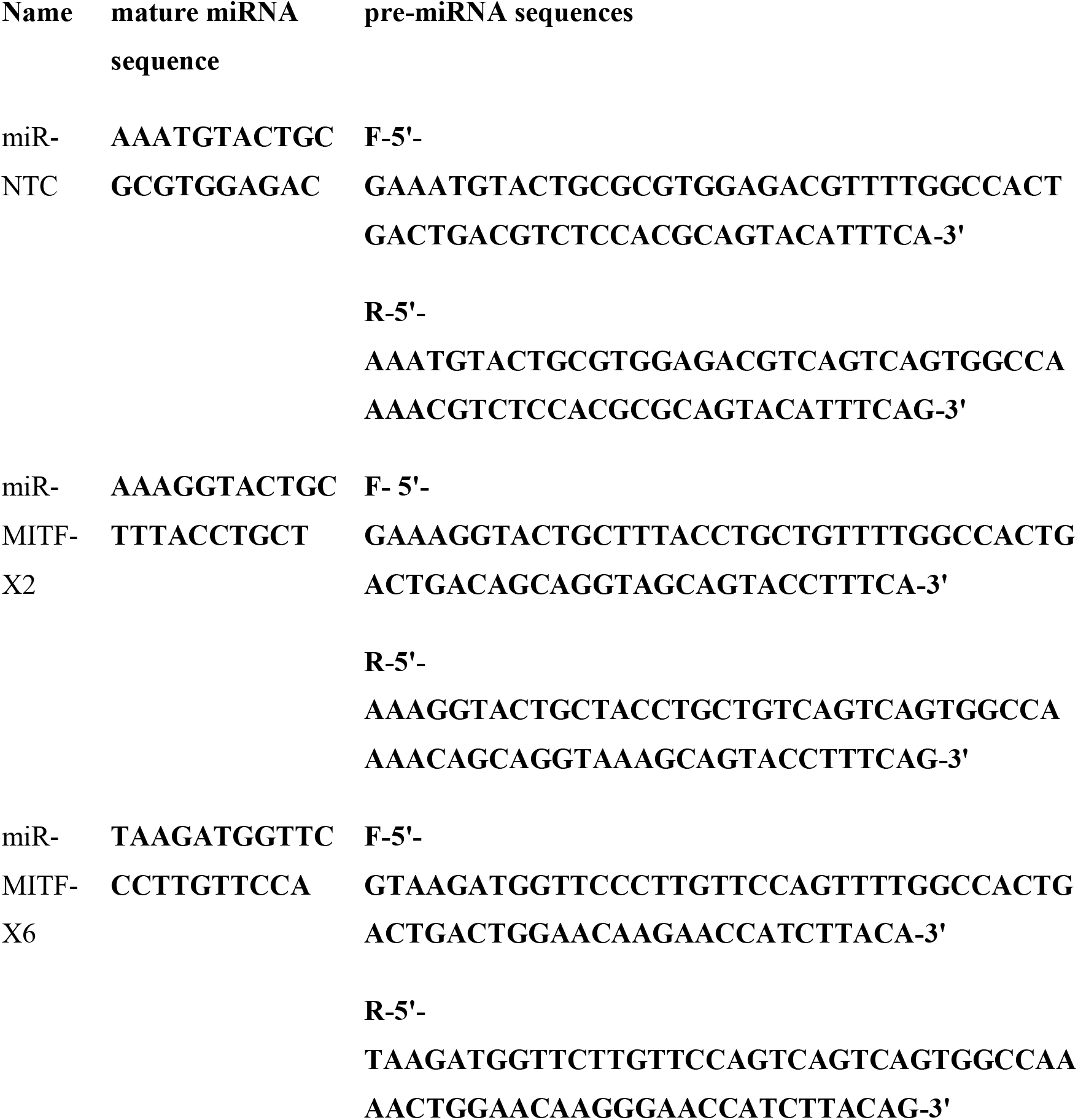
miR sequences used for generating miR-MITF cell lines

For the generation of piggy-bac plasmids containing MITF-M-FLAG-HA and control with only FLAG, we amplified MITF-M cDNA and FLAG sequence from the p3XFLAG-CMV_TM_-14 plasmid expressing mouse Mitf-M using the primers listed in Table 2 (pB-MITF-M-FLAG-HA), and then introduced it into the piggy-bac vector by restriction digestion with *EcoR* I and *Spe* I.

**Table 2.**
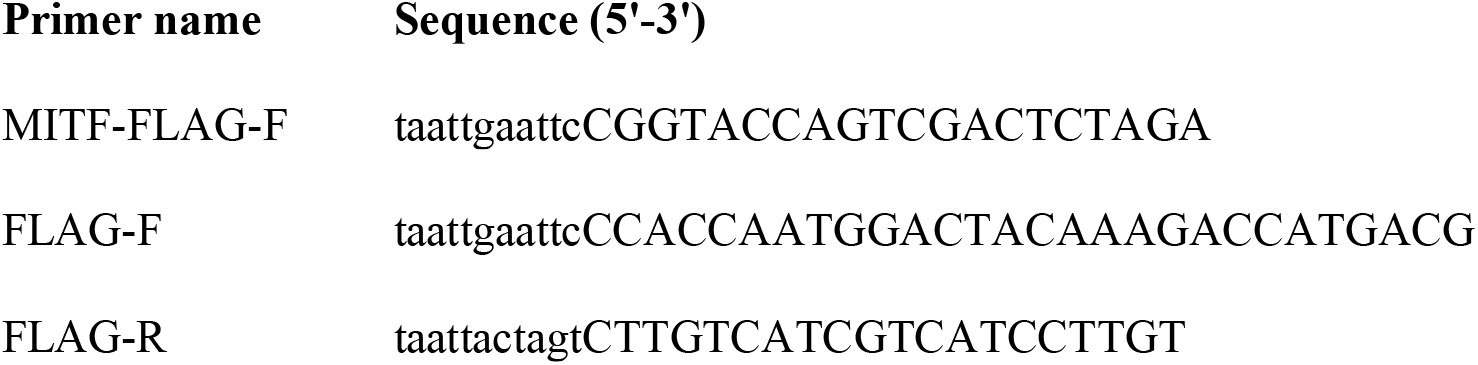
Primers used for generating MITF overexpression lines using piggybac transposon system

### Generation of stable doxycycline-inducible MITF knock down and overexpression cell lines

For generation of stable cells carrying the inducible miR-MITF constructs, 501Mel and SkMel28 cell lines were seeded at 70%-80% confluency and then transfected with the following mixture of constructs: py-CAG-pBase, a vector transiently expressing the piggy-bac transposase, MITF targeting plasmids pBhCMV1-miR(MITF-X2)-pA and pPBhCMV1-miR(MITF-X8)-pA encoding miRNA sequences targeting exons 2 and 8 of MITF, and pPB-CAG-rtTA-IRES-Neo, a plasmid which confers neomycin resistance and rtTA. The mixture was in the ratio of 10:5:5:1, respectively. To generate the miR-NTC controls, 501Mel and SkMel28 cells were transfected at a ratio of 10:10:1 with pA-CAG-pBase, pPBhCMV_1-miR(NTC)-pA encoding a non-targeting miRNA and pPB-CAGrtTA-IRES-Neo. For generation of inducible A375P cells carrying the pB-MITF-M-FLAG or a pB-FLAG empty vector, we transfected 70-80% confluent cells with the following plasmids: py-CAG-pBase, pB-MITF-M-FLAG-HA or pB-FLAG-HA and pPB-CAG-rtTA-IRES-Neo at a 10:10:1 ratio. After 48 hours of transfection, cell lines were subjected to G418 treatment for two weeks (0.5mg/ml, #10131-035, GIBCO) to select for transfected cells.

### RNA isolation, cDNA synthesis and RT-qPCR

Cells were grown in 6-well culture dishes to 70-80% confluency and RNA was isolated with TRIzol reagent (#15596-026, Ambion), DNase I treated using the RNase free DNase kit (#79254, Qiagen) and re-purified with the RNeasy Mini kit (#74204, Qiagen). The cDNA was generated using High-Capacity cDNA Reverse Transcription Kit (#4368814, Applied Biosystems) using 1 μg of RNA. Primers were designed using NCBI primer blast (Table 3) and qRT-PCR was performed using SensiFAST™ SYBR Lo-ROX Kit (#BIO-94020, Bioline) on the BIO-RAD CFX38 Real time PCR machine. The final primer concentration was 0.1μM and 2 ng of cDNA were used per reaction. Quantitative real-time PCR reactions were performed in triplicates and relative gene expression was calculated using the D-ΔΔCt method [65. The geometric mean of β-actin and human ribosomal protein lateral stalk subunit P0 (RPLP0) was used to normalize gene expression of the target genes. Standard curves were made, and the efficiency calculated using the formula E=10[-1/slope].

**Table 3.**
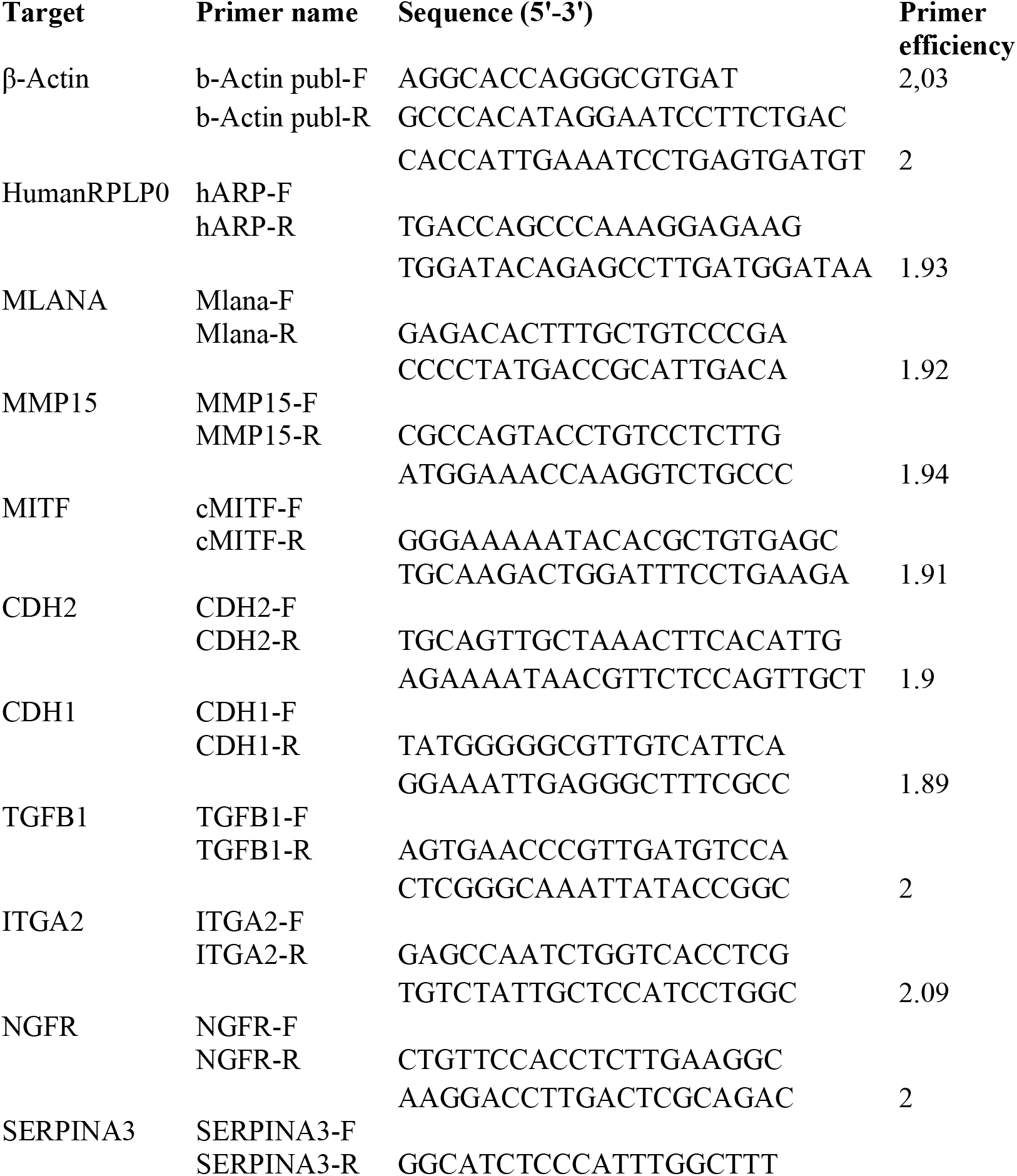
Primers used for RT-qPCR

### Immunostaining

Cells were seeded on 8-well chamber slides (#354108 from Falcon), grown to 70% confluency and then fixed with 4% paraformaldehyde (PFA) diluted in 1xPBS for 15 minutes. After washing 3 times with PBS and blocking with 150 μL blocking buffer (1x PBS + 5% Normal goat serum + 0.3% Triton-X100) for 1 hour at room temperature, cells were stained overnight at 4°C with the appropriate primary antibodies diluted in antibody staining buffer (1xPBS + 1% BSA + 0,3% Triton-X). The wells were washed 3 times with PBS and stained for 1 hour at room temperature with the appropriate secondary antibodies, diluted in antibody staining buffer. The wells were washed once with PBS, followed by DAPI staining at a final concentration of 0.5 μg/ml in 1x PBS (1:5000, #D-1306, Life Technologies) and 2 additional washes with PBS. Subsequently, wells were mounted with Fluoromount-G (Ref 00-495802, ThermoFisher Scientific) and covered with a cover slide. Slides were stored at 4°C in the dark.

### BrdU assay and FACS analysis

Cells were grown on 6-well plates overnight and treated with a final concentration of 10 mM BrdU for 4 hours. The cells were trypsinized and washed with ice cold PBS and then fixed with 70% ethanol overnight. Next, the cells were centrifuged at 500g for 10 minutes and then permeabilized with 2N HCl/Triton X-100 for 30 minutes followed by neutralization with 0.1 M Na2B4O7.10 H2. Cells were analysed on a FACS machine (Attune NxT, Thermo fisher scientific) and data were analysed using FlowJo software.

### IncuCyte live cell imaging

Cells were seeded onto 96-well plates in triplicates supplemented with 200 μL medium with 10% FBS at a density of 2,000 cells per well. Images were recorded with the IncuCyte system at 2 hour intervals for a 4-day period. Images were taken with 10x magnification and four images were collected per well. Collected images were then analysed using the IncuCyte software by measuring cell confluency. Relative confluency was calculated by dividing the confluency at the subsequent hours by the confluency of the initial hour. In addition to this, doubling time was calculated with the following formula: Duration x log(2)/log(FinalConfluency) – log(InitialConfluency).

### Wound scratch and transwell invasion assay

2 x 104 cells were seeded per well of 96-well plate (Nunclon delta surface, Thermo Scientific, #167008) to reach confluent monolayer. Scratches were made with Woundmaker 96 (Essen, Bioscience) and imaging was performed with IncuCyte Live Cell Imaging System (Essen, Bioscience). The recorded images of the scratches were analysed with IncuCyte software to quantify gap closure. For invasion assay transwell chambers with 8μm pore size (Thermo Scientific Nunc™) were coated with matrigel matrix from corning (Thermo Scientific). Then cell suspension of 1×105/300 μl in RPMI 1640 supplemented with 0.1% FBS was added to the matrigel coated upper chamber and the medium containing 10% FBS was added to the lower chamber as a chemoattractant. Cell were allowed to invade for 48 hours after which the cells which migrated to the other side of the membrane were fixed with 4% PFA and stained with DAPI. Images were acquired using QImaging (Pecon, software Micro-Manager 1.4.22) with 10x magnification, and the cells were counted using Image J software.

### RNA sequencing and data analysis

We isolated total RNA as described above from EV-SkMel28 and ΔMITF-X6 cell lines and assessed RNA quality using Bioanalyzer. An RNA integrity (RIN) score above 8 was used for generating RNA libraries. The mRNA was isolated from total 800 ng RNA using NEBNext Poly(A) mRNA isolation module (E7490, NEB). The RNA was fragmented at 94 °C for 16 minutes in a thermal cycler. Purified fragmented mRNA was then used to generate cDNA libraries for sequencing using NEBNext Ultra Directional RNA library Kit (E7420S, NEB) following the protocol provided by the manufacturer with these modifications: Adaptors were freshly diluted 10X before use. A total of 15 PCR cycles were used to amplify the library. A total of 8 RNA libraries were prepared with 4 biological replicates for each cell line including EV-SkMel28 and ΔMITF-X6 cells. Purified RNA sequencing libraries were paired-end sequenced with 30 million reads per sample. Transcript abundance was quantified with Kallisto [66 and index was built with the GRCh38 reference transcriptome. .Differential expression analysis was performed using Sleuth [67 to assess differentially expressed genes between EV-SkMel28 versus ΔMITF-X6. Both likelihood ratio test (LRT) and wald test were used to model differential expression between ΔMITF-X6 and EV-SkMel28 cells. LRT test is more stringent when estimating differentially expressed genes (DEGs), whereas Wald test gives an estimate for log fold change. Therefore, results from LRT test was intersected with Wald test to get significant DEGs with fold change included. We selected differentially expressed genes with the cut off of |log2 (foldchange)| ? 1 and qval <0.05. Functional enrichment analyses (GO terms and KEGG pathway) were performed using Cluster profiler in the Bioconductor R package using Benjamin-Hochberg test with adjusted pvalue <0.05 as a cut-off [68.

Gene set enrichment analysis was performed using GSEA software from the Broad Institute [69. GSEA software was employed with pre-ranked options and gene lists were provided manually to assess enrichment. Differentially expressed genes were ranked combining p-value with log fold change for the input of set enrichment analysis.

### Analysis of human melanoma tumor samples from the Cancer Genome Atlas (TCGA)

The quantified RNA-Seq data from 473 melanoma samples were extracted from the Cancer Genome Atlas database using the TCGAbiolinks package in R Bionconductor [70. The lists of MITF_low_ and MITF_high_ samples were generated by sorting the samples based on MITF expression. The 30 tumour samples with the highest MITF expression and 30 tumour samples with the lowest MITF expression were selected for the downstream differential expression analysis built in the TCGAbiolinks package. Principal Component analysis (PCA) plots were generated using normalized count expression of the 200 most significantly differentially expressed genes between MITF_low_ and MITF_high_ samples and EV-SkMel28 and ΔMITF-X6 cells.

### CUT&RUN

To identify direct MITF target genes, we performed anti-MITF Cleavage Under Targets and Release Using Nuclease (CUT&RUN) sequencing in SkMel28 cell lines as described [71 with minor modifications. Cells in log-phase culture (approximately 80% confluent) were harvested by cell scraping (Corning), centrifuged at 600g (Eppendorf, centrifuge 5424) and washed twice in calcium-free wash-buffer (20 mM HEPES, pH7.5, 150 mM NaCl, 0.5 mM spermidine and protease inhibitor cocktail, cOmplete Mini, EDTA-free Roche). Pre-activated Concanavalin A-coated magnetic beads (Bangs Laboratories, Inc) were added to cell suspensions (200K cells) and tubes were incubated at 4oC for 15 mins. Antibody buffer (wash-buffer with 2mM EDTA and 0.03% digitonin) containing anti-MITF (Sigma, HPA003259) or Rabbit IgG (Millipore, 12-370) was added and cells were incubated overnight at 4oC on rotation. The following day, cells were washed in dig-wash buffer (wash buffer containing 0.03% digitonin) and pAG-MNase was added at a concentration of 500 μg/ mL. The pAG-MNase enzyme was purified in Dr. Robert Cornell’s laboratory following a previously described protocol [72. The pAG-MNase reactions were quenched with 2X Stop buffer (340mM NaCl, 20mM EDTA, 4mM EGTA, 0.05% Digitonin, 100 μg/ mL RNAse A, 50 μg/ mL Glycogen and 2 pg/ mL sonicated yeast spike-in control). Released DNA fragments were Phosphatase K (1μL/mL, Thermo Fisher Scientific) treated for 1 hr at 50oC and purified by phenol/chloroform-extracted and ethanol-precipitated. CUT&RUN experiments were performed in parallel as positive control and fragment sizes analysed using an 2100 Bioanalyzer (Agilent). All CUT&RUN experiments were performed in duplicate.

### Library preparation and data analysis

CUT&RUN libraries were prepared using the KAPA Hyper Prep Kit (Roche). Quality control post-library amplification was conducted using the 2100 Bioanalyzer for fragment analysis. Libraries were pooled to equimolar concentrations and sequenced with paired-end 150 bp reads on an Illumina HiSeq X instrument. Paired-end FastQ files were processed through FastQC [73 for quality control. Reads were trimmed using Trim Galore Version 0.6.3 (Developed by Felix Krueger at the Babraham Institute) and Bowtie2 version 2.1.0 [74 was used to map the reads against the hg19 genome assembly. The mapping parameters were performed as previously described [72. The accession number for the CUT&RUN sequencing data reported in this paper is [GSE153020].

### ChIP-Seq analysis of MITF public dataset

Raw FASTQ files for MITF ChIP-seq were retrieved from GEO archive under the accession numbers GSE50681 and GSE61965 and subsequently mapped to hg19 using bowtie. Peaks were called using MACS, input file was used as control (pval<10e-05) and wig files were generated. Subsequently, wig files were converted to bedgraph using UCSC tool bigWigToBedGraph with the following command line: wigToBigwig file.wig hg19.chrom.sizes output.bw-clip, the hg19, chromosome size file was downloaded from the UCSC genome browser. We used R package ChIPseeker [75 for annotation of ChIP-seq peaks to genes, plotting the distribution of peaks around TSS and a fraction of peaks across the genome. For motif analysis, MEMEChIP [37 was used by extracting DNA sequences corresponding to the peaks that were present in the induced and reduced DEGs of EV-SkMel28 vs. ΔMITF-X6 cells.

### Western blot analysis

200 000 or 100 000 cells were seeded per well of 12 or 6 well cell culture plates overnight and lysed directly with 1X Laemmli buffer (2% SDS, 5% 2-mercaptoethanol, 10% glycerol, 63 mM Tris-HCl, 0.0025% bromophenol blue, pH 6.8), boiled at 95 °C for 10 min and then chilled on ice for 5 minutes. Lysates were spun down for 1 min at 10,000 rpm, run on 8% SDS-polyacrylamide gels and transferred to 0,2 μm PVDF membranes (#88520 from Thermo Scientific). The membranes were blocked with 5% bovine serum albumin (BSA) in Tris-buffered saline/0.1% Tween 20 (TBS-T) for 1 hour at room temperature, and then incubated overnight (O/N) at 4°C with 5% BSA in TBS-T (20 mM Tris, 150 mM NaCl, 0.05% Tween 20) and the appropriate primary antibodies. Membranes were washed with TBS-T and stained for 1 hour at RT with the appropriate secondary antibodies. The secondary antibodies used were the following: Anti-mouse IgG(H+L) DyLight 800 conjugate (1:15000, #5257) and anti-rabbit IgG(H+L) DyLight 680 conjugate (1:15000, #5366) from Cell Signaling Technologies. The images were captured using Odyssey CLx Imager (LICOR Biosciences).

### Statistical analysis

All statistical tests were performed using GraphPad-Prism, one-way or two-way ANOVA was performed and multiple correction was used as indicated in the figure legends.

## Acknowledgements

This work was supported by grants from the Research Fund of Iceland to ES (184861 and 207067) and by a grant from the University of Iceland Doctoral Grants Fund to RD. EEP is funded by the MRC HGU Programme (MC_UU_00007/9), European Research Council (ZF-MEL-CHEMBIO-648489), and L’Oreal-Melanoma Research Alliance (401181). We thank deCODE genetics for their kind assistance with RNA and whole genome sequencing.

---

**Supplementary Figure 1**

(a) Western blot analysis for MITF and Actin in EV-SkMel28, ΔMITF-X2 and ΔMITF-X6 cells. (b) PCR product of exon3, 4, 5 and 6 of MITF in using cDNA generated from 5’RACE experiment.

**Supplementary Figure 2**

(a, b) motif analysis of MITF peaks on reduced and induced DEGs in ΔMITF-X6 vs. EV-SkMel28 cells using MEMEChIP. E-value is a measure of the expected number of motifs with the same size occurring in the random database. (c) View of MITF ChIP-Seq from CUT&RUN in SkMEL28 (rep1 and rep2), MITF ChIP-seq in COLO829 cells [56, and HA-MITF ChIP-Seq in 501Mel [38 loaded in IGV genome browser indicating MITF peaks in *CDH1, CHD2, ZEB1, SNAI2* and *SOX2*.

**Supplementary Figure 3**

(a) Immunostaining for p-PXN_TYR118_ and quantification of p-PXN_TYR118_ positive focal points in EV-SkMel28, ΔMITF-X2 and ΔMITF-X6 cell lines. Error bars represents standard error of the mean, * pval < 0.05 was calculated by one-way ANOVA (multiple correction with Dunnett test) (c) Positive co-expression of MITF mRNA expression and PXN in the 473 melanoma tumour samples displayed in the scatter plot with positive pearson correlation coefficient. (d) Expression of MITF and PXN across 168 melanoma cell lines. Mutations are indicated for each cell line in colours (red:BRAF, orange:NRAS, yellow:cKIT, lime:double, green: WT, and blue: GNA11). (e) Negative correlation of MITF and PXN mRNA expression in 168 melanoma cell lines. (f-h) Immunostaining for p-PXN_TYR118_ and MITF; Quantification of p-PXN_TYR118_ positive focal points in miR-NTC and miR-MITF 501Mel and SkMel28 cells. (i) View of MITF ChIP-seq peaks in *PXN*from ChIP-Seq of CUT&RUN in SkMEL28 (rep1 and rep2), MITF ChIP-seq in COLO829 cells [56, and HA-MITF ChIP-Seq in 501Mel [38 loaded in IGV genome browser.

**Supplementary Figure 4**

(a, b) Western blot analysis and quantification for MITF, ERK and p-ERK inEV-SkMel28, ΔMITF-X2, ΔMITF-X6 and *miR-NTC, miR-MITF* in SkMEL28 and 501Mel cell lines treated with DMSO or vemurrafenib (1μM) for 24 hour. Actin was used as loading control. (c, d) GSEA analysis using DEGs of ΔMITF-X6 vs. EV-SkMel28 cells on Rambow MRD and invasive gene signatures and MRD signature from Zebrafish and mitfalow melanoma tumours. (e) Gene enrichment analysis plotted using Cluster profiler of single cell clusters obtained from melanoma tumours in zebrafish.

**Supplementary Figure 5**

(a,b) Gene expression of *NGFR* and *MLANA* measured by RT-qPCR in EV-SkMel28, ΔMITF-X2 and ΔMITF-X6 cell lines. Expression was normalized to EV-SkMel28 cells. Error bars represent standard error of the mean, * pval < 0.05 was calculated by one-way ANOVA (multiple correction with Dunnett test). (c, d) IGV genome browser showing MITF ChIP-Seq tracks from ChIP-Seq of CUT&RUN in SkMEL28 (rep1 and rep2), MITF ChIP-seq in COLO829 cells [56, and HA-MITF ChIP-Seq in 501Mel [38 in *NGFR* and *MLANA.*

