## Supplementary figures and images for "MITF reprograms the extracellular matrix and focal adhesion in melanoma"

### Supplementary Figure 1

Supplementary Figure 1

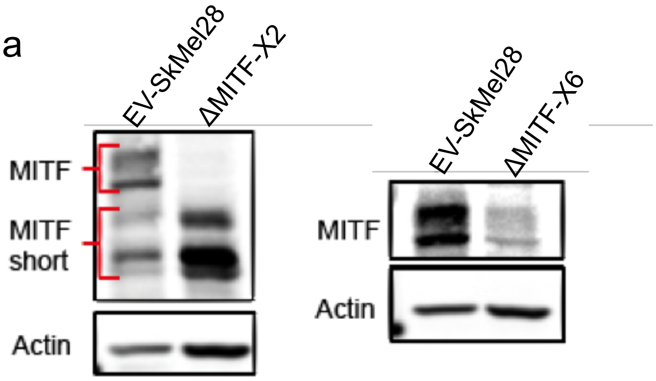


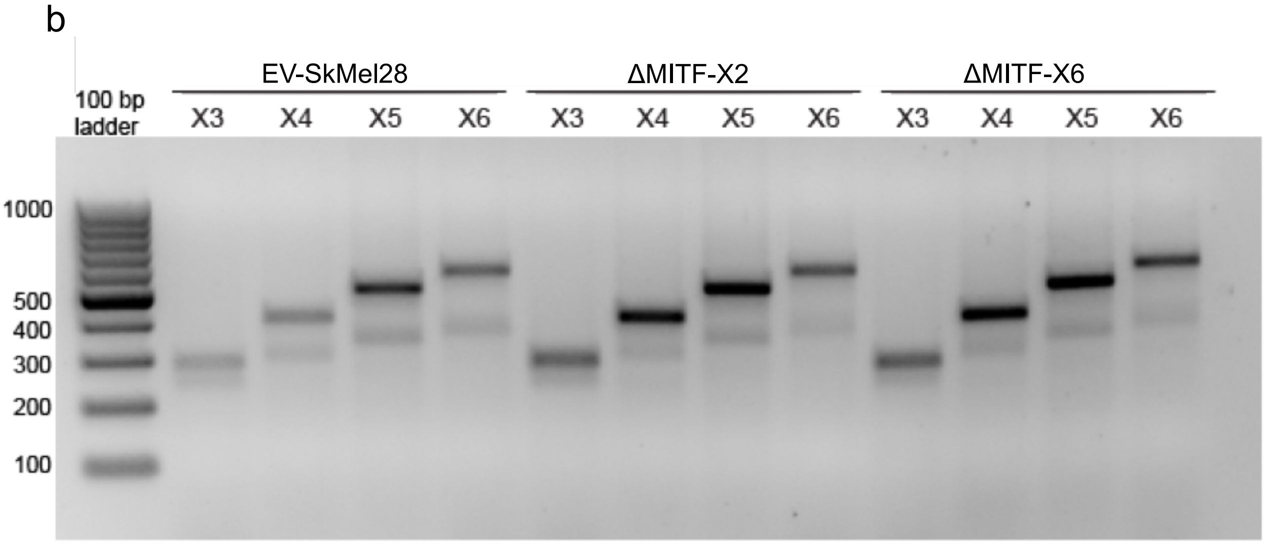

### Supplementary Figure 2

#### Supplementary Figure 2

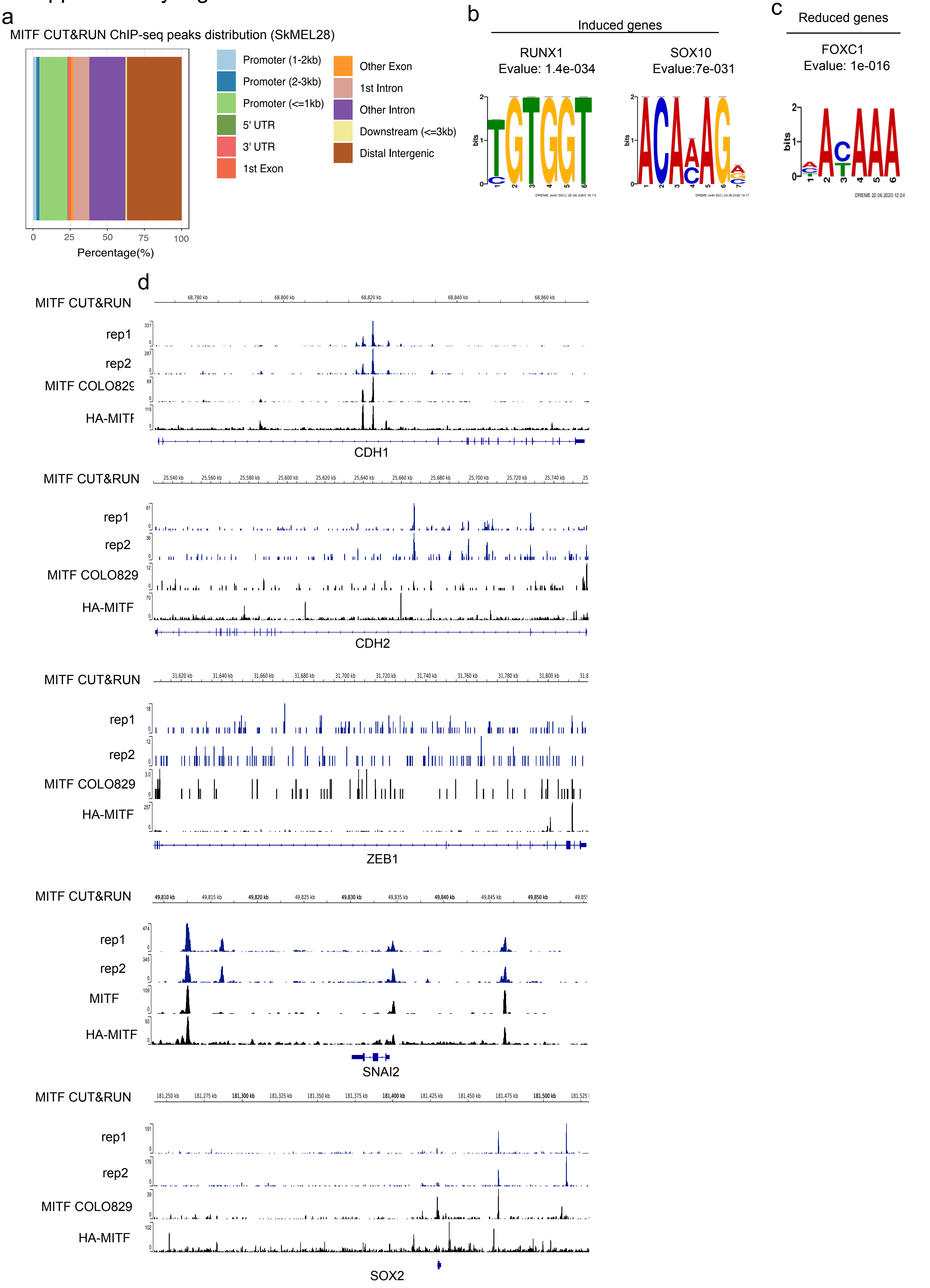

### Supplementary Figure 3

#### Supplementary Figure 3

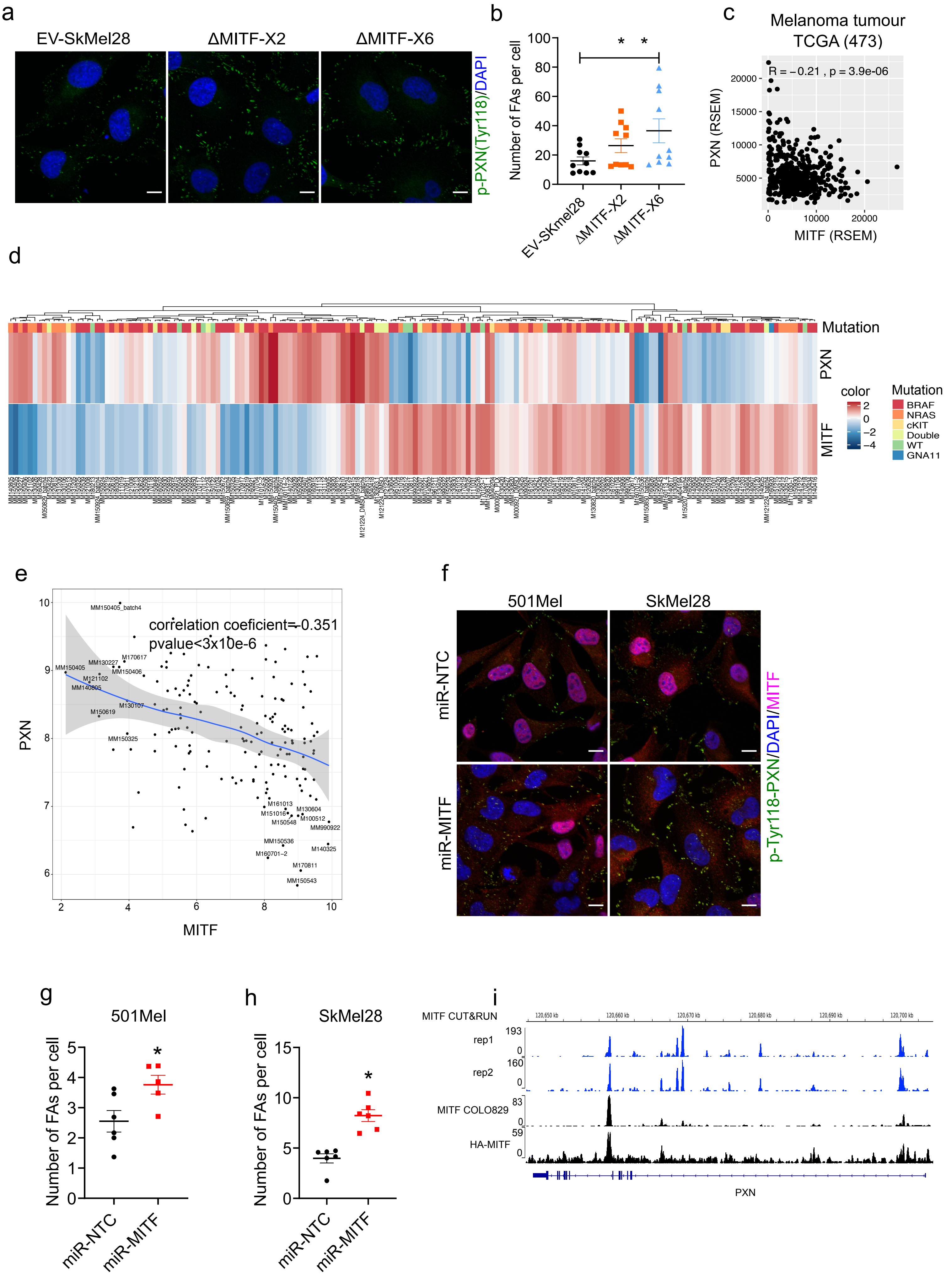

### Supplementary Figure 4

#### Supplementary Figure 4

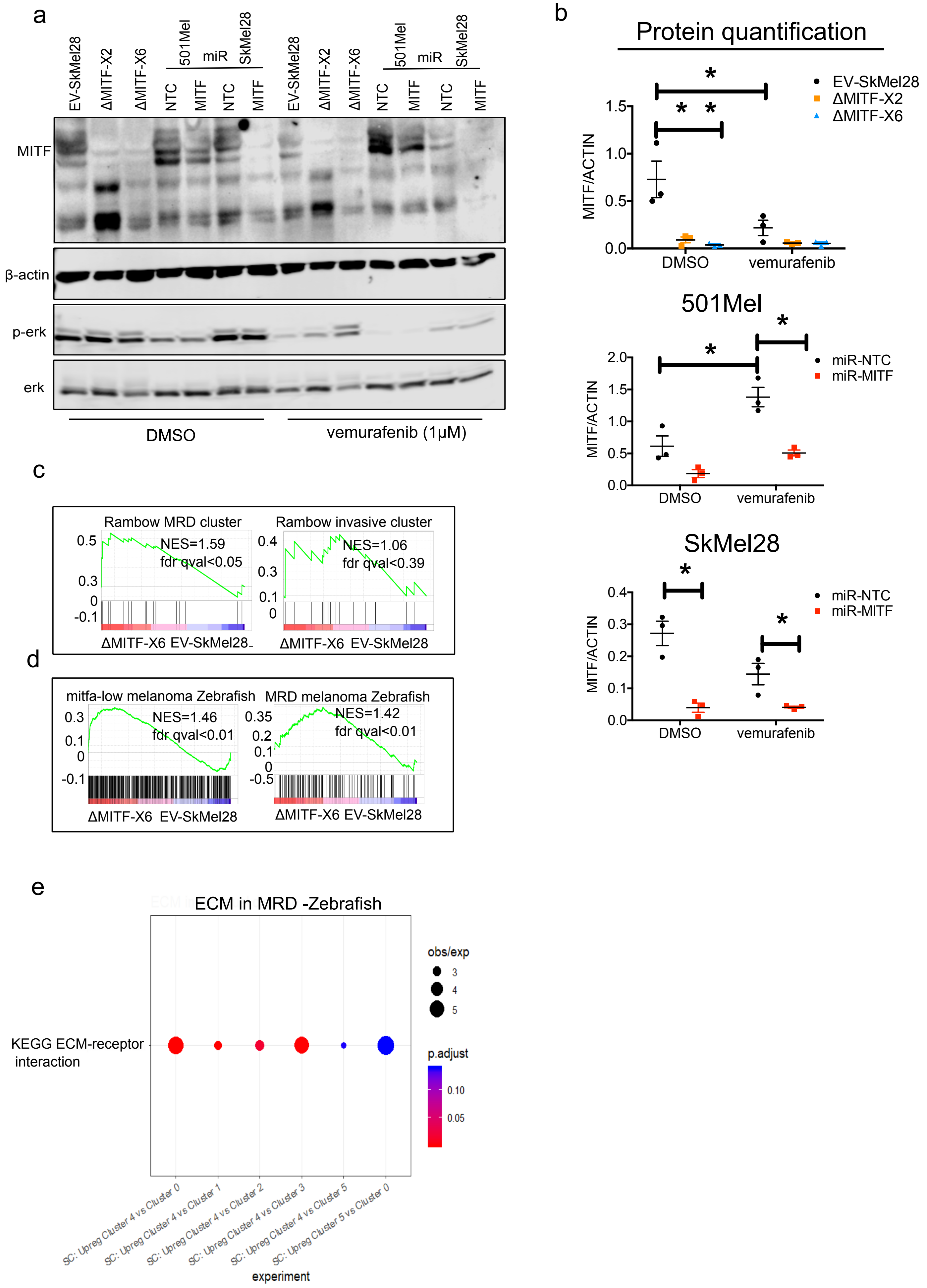

### Supplementary Figure 5

Supplementary Figure 5

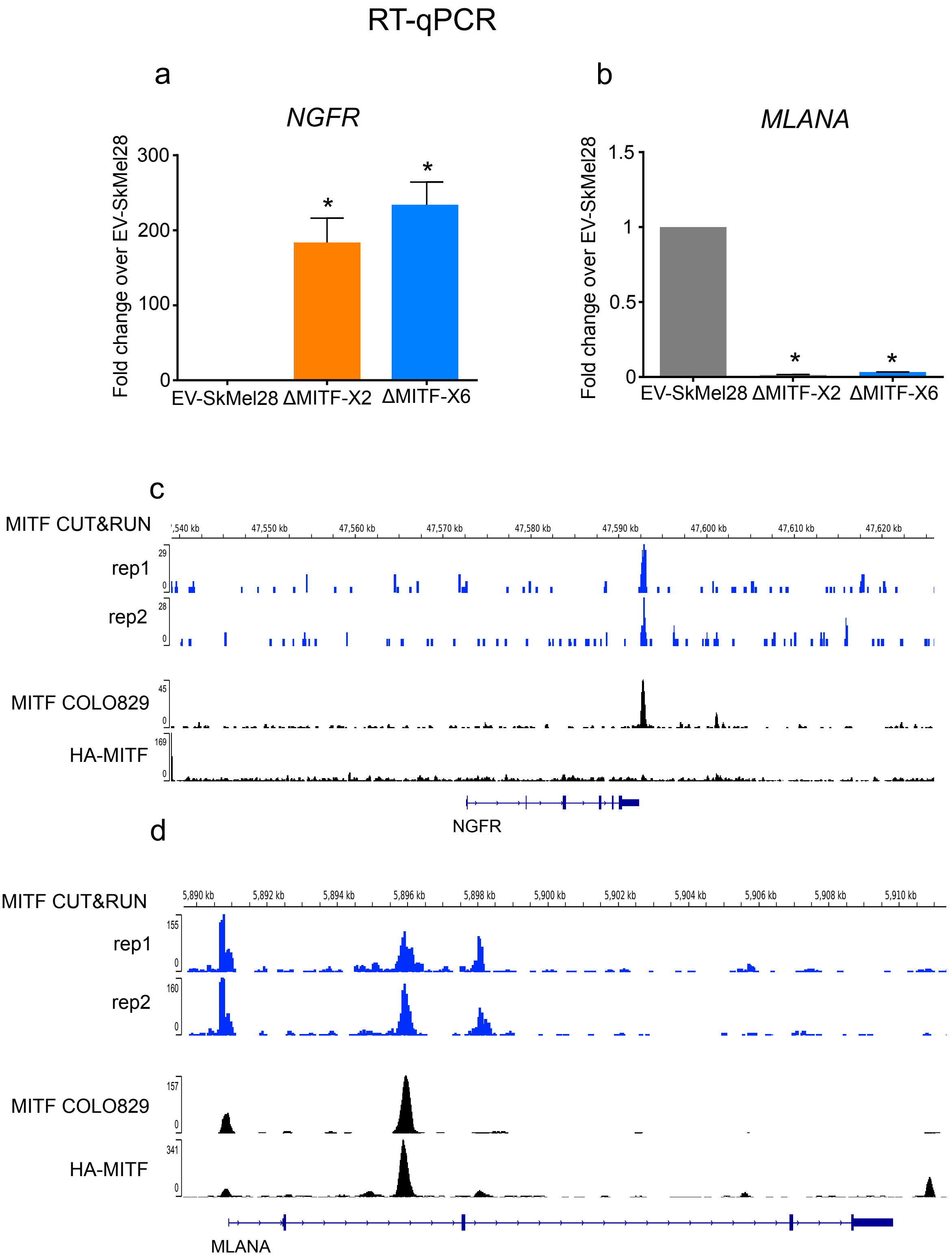
